# Convolutional neural network model to predict causal risk factors that share complex regulatory features

**DOI:** 10.1101/725309

**Authors:** Taeyeop Lee, Min Kyung Sung, Seulkee Lee, Woojin Yang, Jaeho Oh, Seongwon Hwang, Hyo-Jeong Ban, Jung Kyoon Choi

## Abstract

Major progress in disease genetics has been made through genome-wide association studies (GWASs). One of the key tasks for post-GWAS analyses is to identify causal noncoding variants with regulatory function. Here, on the basis of > 2,000 functional features, we developed a convolutional neural network framework for combinatorial, nonlinear modeling of complex patterns shared by risk variants scattered among multiple associated loci. When applied for major psychiatric disorders and autoimmune diseases, neural and immune features, respectively, exhibited high explanatory power while reflecting the pathophysiology of the relevant disease. The predicted causal variants were concentrated in active regulatory regions of relevant cell types and tended to be in physical contact with transcription factors while residing in evolutionarily conserved regions and resulting in expression changes of genes related to the given disease. We demonstrate some examples of novel candidate causal variants and associated genes. Our method is expected to contribute to the identification and functional interpretation of causal noncoding variants in post-GWAS analyses.

## INTRODUCTION

During the last decade, numerous efforts have been made to elucidate the genetic mechanisms underlying complex disorders. Major progress was made through genome-wide association studies (GWASs). However, developing methods to pinpoint the DNA variants that actually increase the risk of the associated disease is a major challenge that GWASs still face (1). GWASs cannot pinpoint causal disease variants but can only report linkage disequilibrium (LD) blocks including many neutral SNPs linked to causal loci. To exacerbate the problem, the majority of disease-associated DNA variations are thought to alter not the gene itself but the regulation of gene expression (2). Our incomplete knowledge of noncoding regions limits the functional interpretation of underlying DNA variants. Fortunately, the wealth of cell-type-specific human epigenomes help with the identification of functional noncoding variants (1, 3–5).

Deep learning is a powerful approach for learning complex patterns (6) and has been applied in genomics especially for deciphering the complexity of noncoding DNA sequences. For example, a deep-learning model that predicts regulatory codes for RNA splicing has been developed (7). As one of the most successful deep learning architectures, convolutional neural networks (CNNs) have been used to systematically learn the sequence motifs or regulatory patterns embedded in genomic regions recognized by DNA- or RNA-binding proteins (8) or in DNase I hypersensitive sites (DHSs) (9).

In this study, we developed a deep learning framework based on CNNs to discover regulatory variants that may play a causative role in increasing the risk of the five major psychiatric disorders and four autoimmune diseases: autism spectrum disorder (ASD), attention deficit-hyperactivity disorder (ADHD), bipolar disorder (BPD), major depressive disorder (MDD), schizophrenia (SCZ), rheumatoid arthritis (RA), systemic lupus erythematosus (SLE), Crohn’s disease (CD), and ulcerative colitis (UC). We utilized numerous functional features while combining them nonlinearly to model complex patterns shared by risk variants. We were able to discover novel candidate SNPs that may actually contribute to disease development.

## MATERIALS AND METHODS

### Identification of association blocks

In order to make the input data for the CNN model, we first identified chromosomal association blocks. To this end, we employed a cross-disorder GWAS dataset for five major psychiatric disorders and four autoimmune diseases (Supplementary Table 1) (10, 11). The association P values were downloaded from the Psychiatric Genomic Consortium portal (https://www.med.unc.edu/pgc/results-and-downloads) for psychiatric disorders. As for the autoimmune diseases, the association P values were retrieved from the largest meta-analysis for each disease (12–14). For imputation, we used the EUR (European ancestry) samples of the 1000 Genomes Project phase 1 release (version 3) as the reference panel (15). Imputation of summary statistics was performed by using ImpG-Summary (16). The SNPs that showed the strongest associations and were at least 1 Mb apart from one another were defined as lead SNPs. To identify the lead SNPs, we sorted all SNPs across the whole genome according to the observed or imputed P values and picked SNPs from the top of the list while maintaining a > 1 Mb distance from each of the previously selected ones. We then searched upstream and downstream regions flanking each lead SNP for the 30 most significant SNPs. We then discarded those with P > 5×10^-4^. In this manner, we identified association blocks carrying the lead SNP and their neighboring SNPs while constraining the maximum number of neighboring SNPs to 30 (Figure 1a). The resulting number of association blocks was 340, 391, 474, 405, and 601 in ADHD, ASD, BPD, MDD and SCZ for psychiatric disorders, and 435, 849, 431, and 383 in RA, SLE, CD, and UC for autoimmune diseases, respectively (Supplementary Table 2).

**Figure 1.**
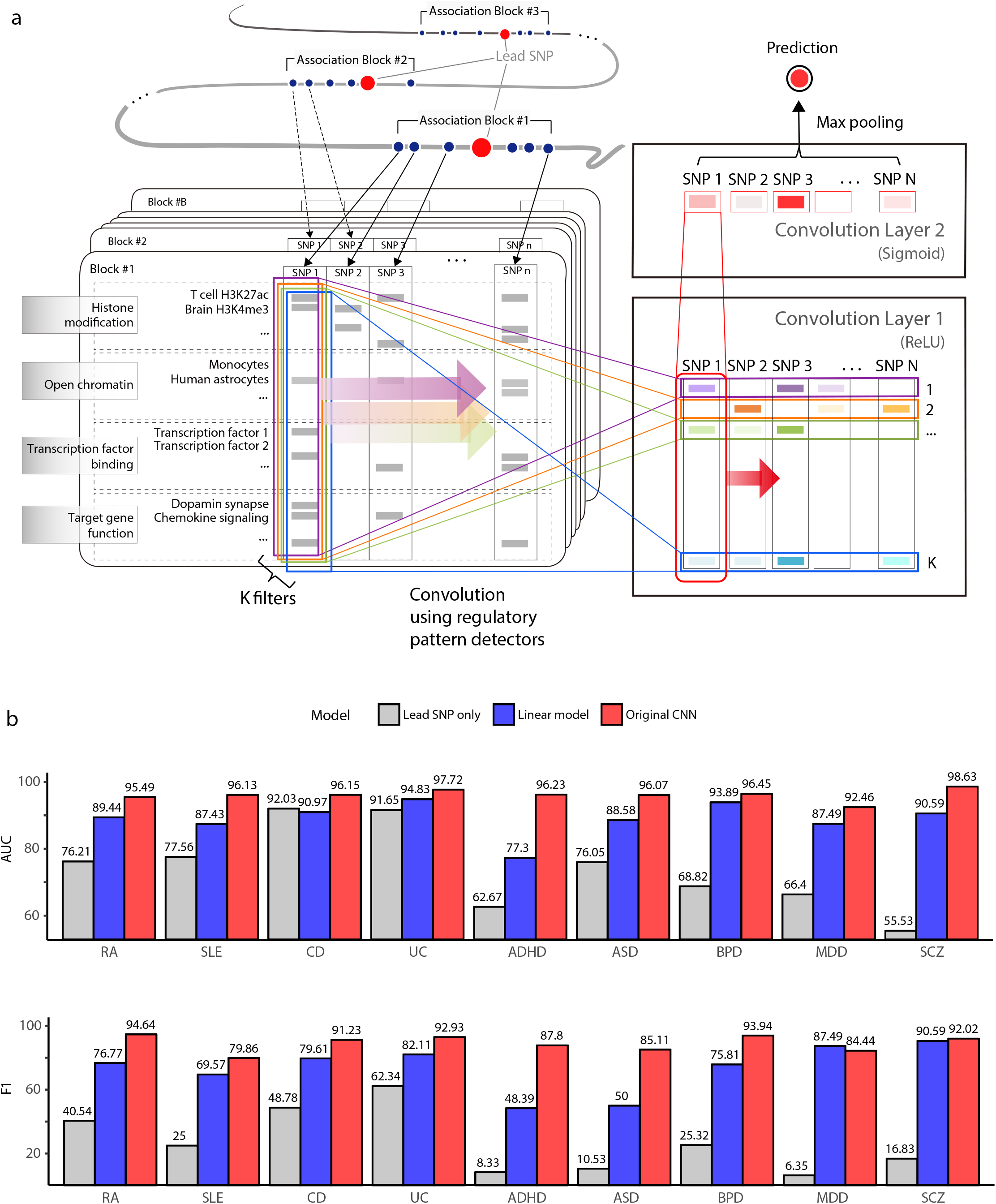
Model schematic and performance. **(a)** CNN framework to detect regulatory patterns shared by risk variants residing in multiple association blocks centered on lead SNPs. In this example, block #1 carries *n* SNPs including the lead SNP. We apply *k* different kernels that learn particular patterns composed of various regulatory features encompassing DHSs, histone modifications, target gene function, and TF binding sites. At this stage, an autoencoder is used for pre-training. In this manner, the first convolution layer scores *n* SNPs with *k* pattern detectors. Afterward, another convolution layer is applied to combine the *k* scores, thereby enabling nonlinear combinatorial modeling of regulatory patterns. The output of the second layer serves as the prediction score for each SNP. The model is trained to maximize the likelihood derived from the block scores that are assigned by max pooling. **(b)** Model performance of rheumatoid arthritis (RA), systemic lupus erythematosus (SLE), Crohn’s disease (CD), ulcerative colitis (UC), attention deficit-hyperactivity disorder (ADHD), autism spectrum disorder (ASD), bipolar disorder (BPD), major depressive disorder (MDD) and schizophrenia (SCZ) measured on the basis of AUC and F1. The red, blue, and gray bars are for the original CNN model, linear model with only one convolution layer, and model with only the lead SNPs. Model training and performance evaluation were carried out on the training, validation, and testing sets (Supplementary Figure 2 and Supplementary Table 2).

### Feature set construction

The overall processes for our feature map construction are illustrated in Supplementary Figure 1. We obtained open chromatin profiles in 349 different samples as provided in the form of DHS peaks by the ENCODE Project (17) and the Roadmap Epigenomics Project (18). A total of 606 histone modification profiles from the Roadmap Epigenomics Project (18) were obtained as the narrow peak bed files in 127 different biological conditions. The KEGG database (20) was utilized to incorporate gene functions as features. Each SNP was mapped to their target gene and its KEGG pathway when they resided in the gene body or 500 kb upstream of the TSS. We ran FIMO (http://meme-suite.org/doc/fimo.html) (21) to search the TRANSFAC (22) and JASPAR (23) databases of position weight matrices for 996 transcription factors (TFs). We used the P value threshold of 10^-4^ for feature mapping. As a result, we compiled 2,252 functional features consisting of DHSs, histone modifications, KEGG pathways, and TF binding sites. Because using too many features results in overfitting, we filtered features that were shared by only a small number of SNPs in the association blocks (Supplementary Figure 1). Specifically, we retained the features that were mapped to any SNPs in > 95% of the association blocks in each disease model. The resulting number of features was 714 for ADHD, 711 for ASD, 730 for BPD, 714 for MDD, 739 for SCZ, 791 for RA, 741 for SLE, 845 for CD, and 834 for UC.

### Model design

The developed CNN model consists of hierarchical pattern detectors that learn the regulatory features commonly present in disease-associated genomic loci. In our model, we used two convolution layers; 1) the first layer acts as a local feature extractor at the individual SNP level, and 2) the second layer combines the detected patterns into more complex biological features.

#### 1) Convolution layer 1

The convolution layer 1 takes an input matrix (*X_mn_*) with a size of *m* × *n* as described below:

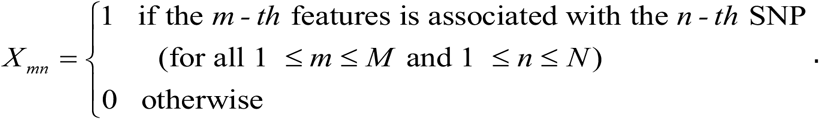

*M* is the number of regulatory features that survived the filtering step described above, and *N* is the total number of candidate SNPs in an association block. With this input matrix and tunable patterns represented as kernel *w^k^*, the first feature map array 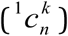 was obtained by the convolution process,

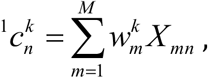

where *k* is the index for 50 filters used in our model (1 ≤ *k* ≤ *K, K* = 50). Each convolution kernel *w^k^* is a weight vector with a length of *M*. This process is equivalent to performing a one-dimensional convolution (8) with a moving window of step size 1 on the list of consecutive *N* SNPs each having *M* channels. As implied in the above formula, *K* types of pattern detectors were used for each SNP without considering the effect of neighboring SNPs. After convolving the matrix *X_mn_*, we added a bias vector, *b_k_*, and then applied a rectified linear unit (ReLU) as follows:

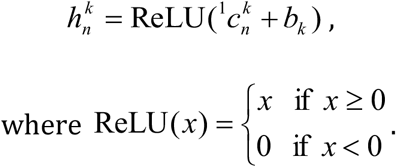

We then stacked the output vector 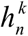 to compose a matrix *H_kn_* with a size of *k* × *n*, which corresponds to scores measuring how well the features of each SNP match the patterns of the shared weights.

#### 2) Convolution layer 2

The convolution layer 2 operates on the output matrix of the previous layer (*H_kn_*). The second feature map vector (^2^*c_n_*) was obtained as

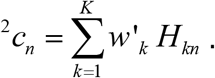

In this step, only one tunable weight vector was used to linearly combine the *K* patterns for high-level feature scoring of each SNP. Then, we added a bias term *b′* and scaled ^2^*c_n_* + *b′* to the 0-1 range by the sigmoid function:

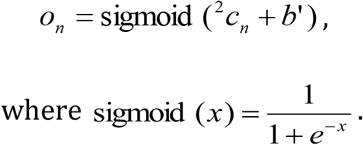

*o_n_* can be regarded as the prediction score of the *n*-th SNP. Prediction scores close to 1 indicate that certain common regulatory patterns are embedded in the features of the given SNPs. Finally, we applied max-pooling by taking the maximum of *o_n_* as

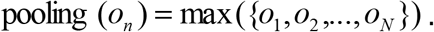

This corresponds to the per-block score of the SNP whose features best match the common patterns shared by different association blocks.

### Model training

The input data for the model was split into training, validation, and testing sets (Supplementary Figure 2 and Supplementary Table 2). The validation and testing sets were used to select the best hyperparameter set and report final performance levels, respectively. We trained the model parameters to minimize negative log likelihood (NLL) defined as follows:

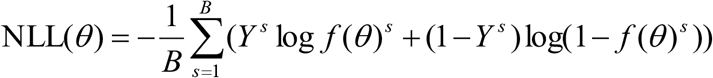

where *f*(*θ*)^*s*^ is an output for the *s*-th block in a mini-batch of size *B* from the training set (*B* =100 was used for all experiments). Each target, *Y*, serves as the dependent variable and can be either 1 (for true cases or the blocks that carry the pattern of functional SNPs associated with the disease) or 0 (for false cases or the blocks that are unlikely to carry the pattern of functional SNPs associated with the disease). We constructed false cases by shuffling the regulatory features of SNPs in the true cases. In doing so, it is anticipated that the features of false cases do not reflect any disease-related common patterns. The shuffling procedure was done separately for each feature group. We generated false cases that are ten times the true cases. The NLL was composed of parameters (*θ*) including 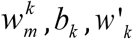 and *b*′, which were updated by the standard backpropagation algorithm with momentum. To prevent overfitting, early stopping and pre-training by an autoencoder were used. More details of the model training processes are documented in **Supplementary Information**.

### Feature importance analysis

Using the model weights learned by CNN for the assessment of feature importance can be misleading. To tackle this drawback, we employed random forest to assess the relative importance of each feature (24). The input SNPs for training RF were labeled as positive (prediction score > 0.5) or negative (prediction score < 0.5) according to our CNN results. For each feature, we calculated the Mean Decrease Gini (MDG), which is defined as the average of total decrease in Gini impurities in each tree. A greater MDG indicates higher importance of a feature. The empirical P value of MDG was calculated by 1,000 random permutations. The hypergeometric distribution was used to test whether disease-related features stood out in terms of the MDG. Neural and immune features were defined on the basis of open chromatin or histone marks in neuronal cells or tissues and in immune-related blood tissues (all lymphoid cells and granulocytes) or cell lines, respectively. KEGG pathway terms under categories “nervous system”, “neurodegenerative disease” and “substance dependence” were included as neural features, while “inflammatory process”, “immune process” and “chemokine signaling pathway” were included as immune features. As negative controls, we used irrelevant features consisting of open chromatin peaks, histone modifications, and KEGG pathways related to the digestive and circulatory system in the same way. In our network analysis, the neural features with a significant (P < 0.05) MDG were selected for visualization. RF model building and feature importance analysis were implemented by using R packages randomForest and rfPermute (25). Network visualization of the significant neural features was based on Cytoscape (v3.5.1) (26).

### Functional analyses of predicted variants

Details on the TF binding and allelic imbalance analysis, evolutionary conservation analysis, and target gene function analysis are provided in **Supplementary Information**. Additional dataset not used in the training process was employed for the functional analysis of the autoimmune diseases. A total of 100 histone modification profiles in 16 blood tissues or cell lines were obtained as peak bed files from the BLUEPRINT epigenome project (www.blueprint-epigenome.eu) (19).

## RESULTS

### Prediction model and performance

Our CNN model was trained on the feature vectors across multiple association blocks (Figure 1a and Supplementary Figure 1). The dependent variable of the model is 1 for true cases or the blocks that carry the pattern of functional SNPs associated with the disease and 0 for false cases or the blocks that do not carry the pattern of functional SNPs associated with the disease. The premise of the model is that there are one or more functional variants in association blocks, and that many of the functional variants share certain patterns of regulatory features despite being scattered in different blocks. Therefore, the association blocks identified through GWASs served as true cases.

In analogy, a true case (association block) can be compared to a face image, and functional SNPs can be compared to eyes, nose, or mouth. By observing many face images, a CNN model can learn that a face has eyes, nose, and mouth in common, and decide whether a given picture is a face or not. Likewise, if a CNN model is trained with multiple true cases, it can learn that association blocks carry functional SNPs with certain patterns and can decide whether a given region is a true case.

The number of association blocks used as true cases for ADHD, ASD, BPD, MDD, SCZ, RA, SLE, CD, and UC was 340, 391, 474, 405, 601, 435, 849, 431, and 383, respectively (Supplementary Table 2). These blocks were partitioned into the training set, validation set, and testing set (Supplementary Table 2). Details of the learning processes are summarized in Supplementary Figure 2. Performance evaluation was based on the area under the receiver-operator characteristic curve (AUC) and F1 value (Figure 1b). We modified our model to learn the features of only the lead SNPs (i.e., the most significant SNPs indexing each association block) or to learn patterns composed of the linear combinations of features (by using only one convolution layer). The lowered performance of the modified models (gray and blue bars of Figure 1b) indicates that common regulatory patterns need to be searched for through all variants in each chromosomal block in a complex, non-linear fashion. This justified the usage of a CNN model for this task.

Overall, the lowest performance was achieved for MDD, probably reflecting that genetic factors play a less significant role relative to the other diseases (27). We defined positive calls as variants that were assigned a prediction score greater than 0.5. The list of these putatively causal variants in each disease is provided in Supplementary Table 3.

### Biological validation of prediction results

First, true causal variants are expected to have a certain level of statistical association with the given phenotype. Indeed, our model assigned higher prediction scores to associated variants in the testing set, which is independent of the training processes (Supplementary Figure 3). In >50% of the chromosomal blocks with at least one positive call, the variant with the strongest statistical association (i.e., lead SNP) was positively predicted (Supplementary Table 2). Also, there were many cases in which the greatest prediction score was assigned to the lead SNPs (Supplementary Figure 4). Of importance, our functional prediction method was able to single out one of statistically indistinguishable variants (compare red diamonds and blue circles in Figure 2).

**Figure 2.**
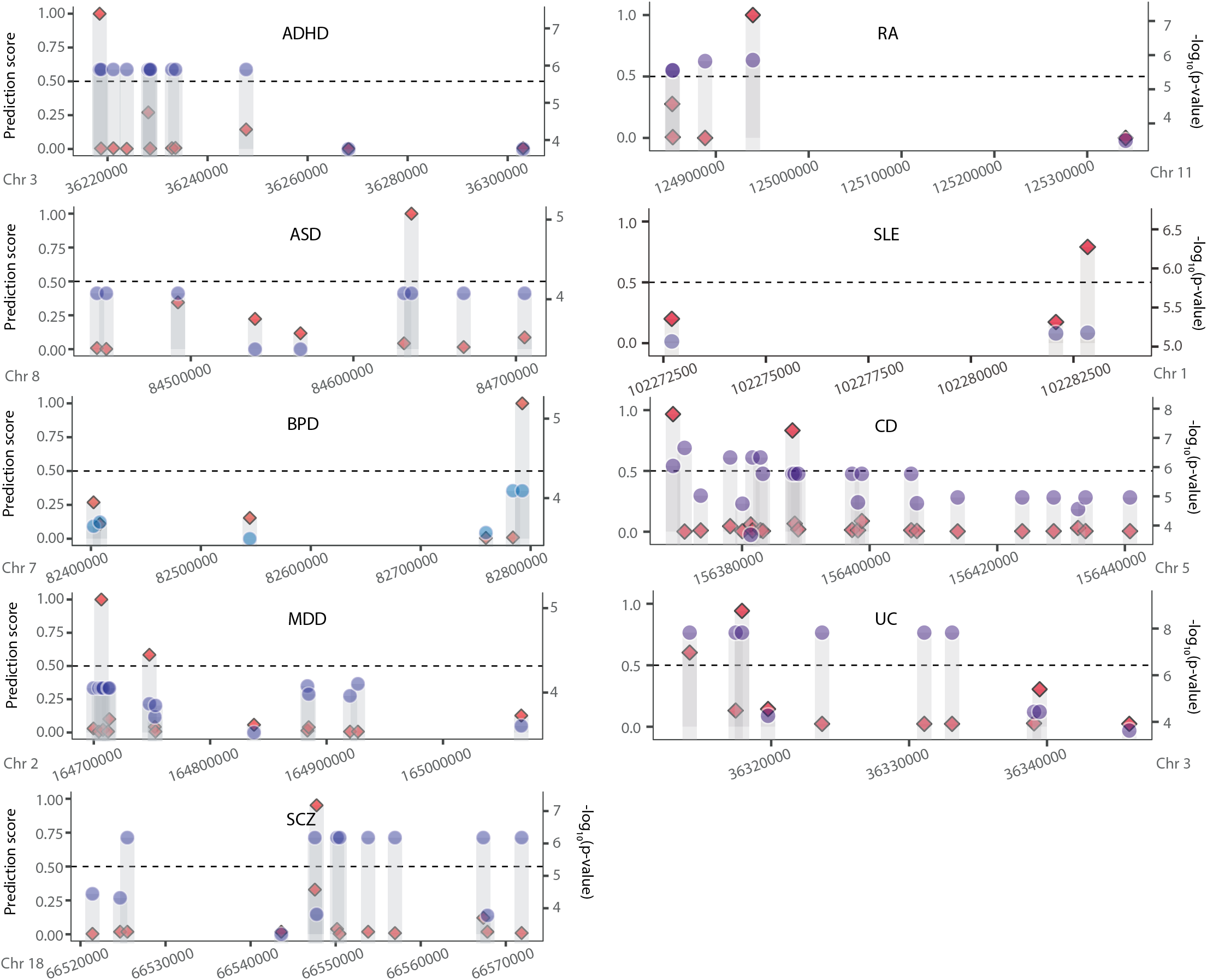
Comparison of our prediction scores (red diamonds on the left y-axis) and association statistics (blue circles on the right y-axis) for individual SNPs in exemplary association blocks. Our functional prediction enables to discern statistically indistinguishable variants.

Second, disease-related features are expected to play an instrumental role when predicting causal variants. For example, psychiatric disorders should be associated with brain-related features while autoimmune diseases with immune-related features. To test this, we employed the random forest classifier to assess the contribution of each feature to the prediction processes. By randomizing each feature, the explanatory power of the given feature in discriminating positive and negative calls could be estimated. We used the MDG score for this measure (see Materials and Methods). With this metric, we observed higher discriminative power for neural features and immune features than irrelevant features in the psychiatric disorders and autoimmune diseases, respectively (Figure 3a). A comprehensive target gene analysis further supported the clinical relevance of our prediction (Supplementary Figure 5).

**Figure 3.**
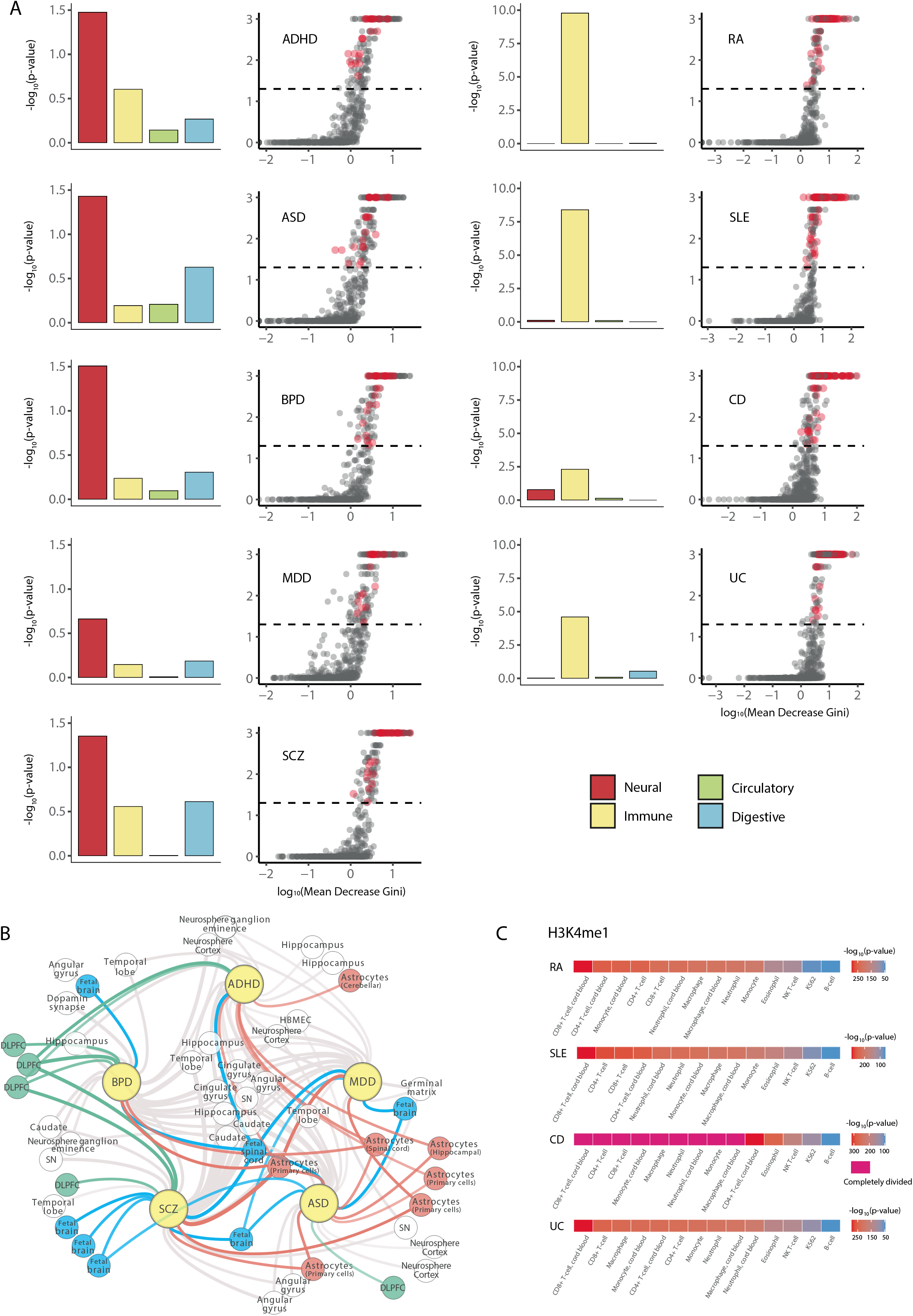
Pathophysiological relevance of prediction outcomes. **(a)** Results of feature importance analysis. The MDG score (x-axis) and its P value (y-axis) on the log scale with significant (P < 0.05) neural features highlighted in red (left panel). Enrichment of feature categories estimated by the hypergeometric test (right panel). **(b)** Network of neural features that were important in each disease model. The small nodes represent neural features that showed a significant (P < 0.05) MDG in the prediction model of the connected disease. The significant neural features, including those related to fetal brain (blue), astrocytes (red), and dorsolateral prefrontal cortex (green), were mapped to the relevant disorder (yellow). **(c)** Enrichment of positive calls for autoimmune diseases for the BLUEPRINT epigenome data for various immune cell types. Positive calls are more enriched in the regulatory regions of lymphocyte lineages rather than granulocytes. Shown here are regulatory regions marked by H3K4me1. The H3K27ac data is provided in Supplementary Figure 6.

In addition, some neural features reflected the pathophysiology of the relevant psychiatric disorder (Figure 3b). For example, astrocyte (red nodes) and dorsolateral prefrontal cortex (green nodes) are often implicated in ASD (and ADHD) and SCZ (and BPD), respectively. Fetal features (blue nodes) were important when characterizing neurodevelopmental disorders such as ASD and SCZ. Similarly, the BLUEPRINT epigenome data for various immune cell types were used for the analysis of the autoimmune disease results. Positive calls were more enriched in the regulatory regions of lymphocyte lineages rather than granulocytes. This is in good agreement with the pathophysiology of autoimmune diseases (Figure 3c & Supplementary Figure 6).

Third, causal noncoding variants are likely to lie in the regulatory regions of relevant tissue types. We first tested whether positive calls are enriched in regulatory regions derived from independent data. The BLUEPRINT epigenome data for various immune cell types were useful for this purpose because they were not used for model training. The fractions of positive calls for the autoimmune diseases were significantly higher than negative calls in H3K4me1 and H3K27ac regions from the BLUEPRINT epigenome data (Figure 4a). Next, to compare disease-relevant tissues with others, we ordered all available tissue types from the Epigenome Roadmap project depending on the degree of positive call enrichment. Expectedly, positive calls were enriched in the disease-related tissues for both the psychiatric disorders and autoimmune diseases (Figure 4b).

**Figure 4.**
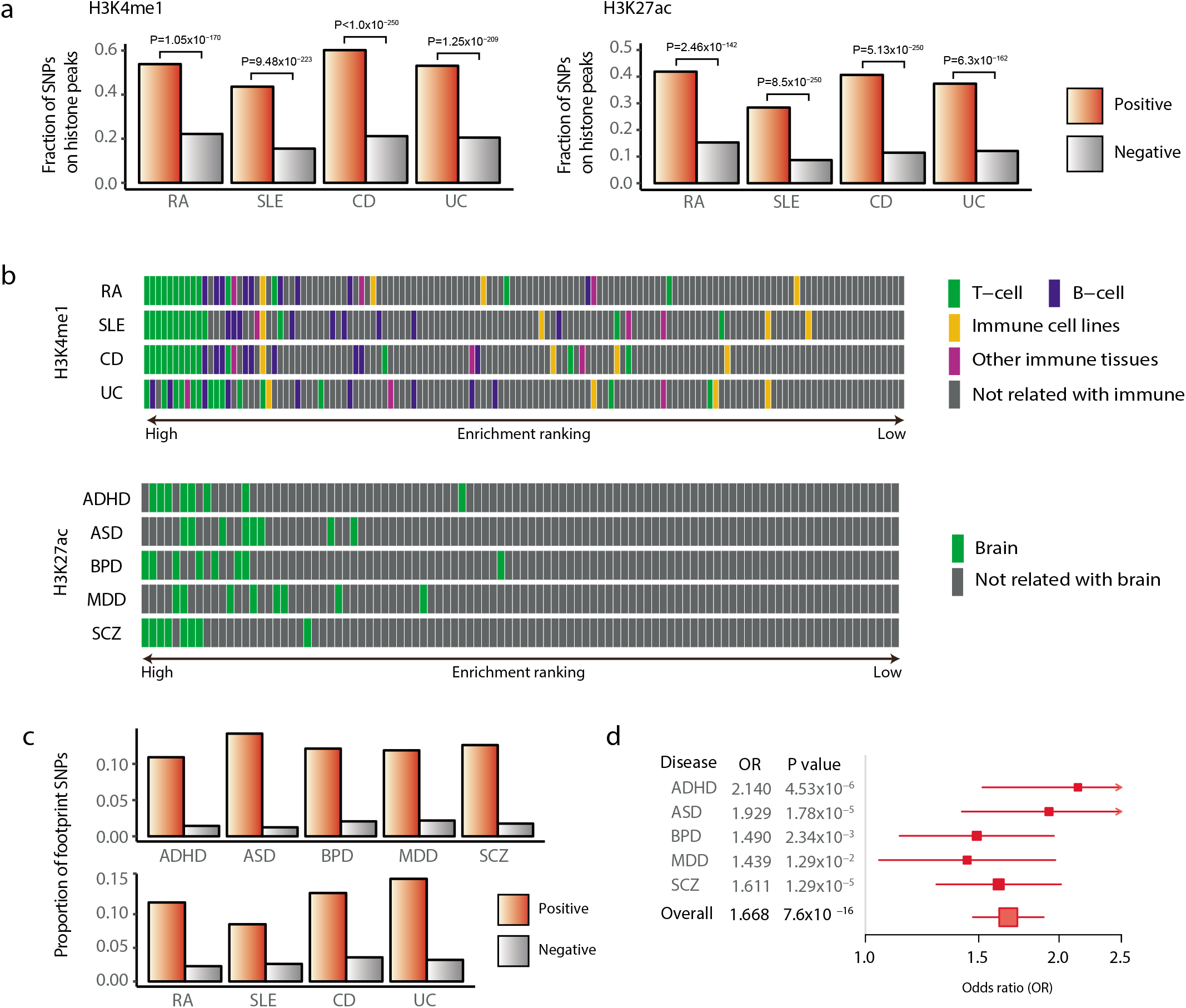
Functional relevance of prediction outcomes. **(a)** Fractions of positive calls and negative calls for autoimmune diseases in H3K4me1 and H3K27ac regions from the BLUEPRINT epigenome data. **(b)** Enrichment of positive calls for autoimmune diseases in immune-related cells and non-immune cells from the Epigenome Roadmap project. **(c)** Proportion of positive (prediction score >0.5) SNPs and negative (prediction score < 0.5) SNPs that match TF footprints. **(d)** The ratio of the odds of positive calls in conserved regions to their odds in non-conserved regions. For the conserved regions, we searched association blocks for the primate PhastCons score > 0.5. The odds of positive calls were computed as the ratio of the positive to negative SNPs in the conserved or nonconserved regions. Shown is the odds ratio together with its 95% confidence interval and P value.

Fourth, the mechanisms by which noncoding variants contribute to disease phenotypes should involve TF binding changes. For this test, we utilized TF footprint data generated by base-resolution DHS analyses (28). The positive hits included a significantly higher fraction of nucleotides that are in contact with cognate TFs than the negative calls (Figure 4c). A useful method to test the functionality of regulatory variants is to examine allelic imbalance in chromatin accessibility (29). By examining allelic patterns in footprint reads, we found that different alleles at positive calls are more likely to cause distinct regulatory variation (Supplementary Figure 7a). We also tested whether the predicted causal variants tend to affect gene expression levels. When examined using the 1000 Genomes whole-genome and transcriptome data, positive calls showed higher levels of expression association, further supporting the functionality of the predicted variants at the transcription level (Supplementary Figure 7b).

Finally, true causal variants for major psychiatric disorders are likely to reside in regions that are critical for brain development and function. Therefore, one can anticipate higher evolutionary conservation, especially among primates, for regions surrounding the causal variants. Indeed, the odds of positive calls in conserved sequences were statistically significant (Figure 4d). This tendency was more distinct with ASD, ADHD, and SCZ, in which aberrations in neural development play a critical role, as compared to MDD and BPD. Also, the average degree of sequence conservation was markedly higher for genomic regions centered on positive calls than negative calls (Supplementary Figure 8a). The degree of conservation was less significant when measured among mammals or vertebrates (Supplementary Figure 8b).

### Examples of novel candidate causal variants

As shown in Figure 1b, our method was able to detect common patterns shared by variants other than the lead SNPs of association blocks. In other words, there must be numerous cases in which the tag SNPs or lead SNPs detected by the typical GWAS analysis based on statistical association may not act as causal variants for the disease. For example, a known GWAS variant of RA (30), rs773125, located on chromosome 12, has the strongest association with the RA phenotype (Figure 5a). Previous GWAS studies assigned this SNP to CDK2 on the basis of physical distance. However, the nearest gene is not always the actual target gene. Only a small fraction of distal enhancers target the nearest transcript (31). Not surprisingly, CDK2 has no clear role in association with RA. Furthermore, rs773125 is not located in active regulatory regions marked by H3K4me1 or H3K27ac in any types of immune-related tissues. Our prediction was negative on rs773125. Instead, there were two positive SNPs (rs773114 and rs1873914) that showed a lower association with RA. GM12878 capture Hi-C data (32) located these SNPs in an enhancer region of RPS26 (Figure 5a). This site was an active regulatory region of CD14+ monocyte as well. There were several studies that implicated RPS26 in autoimmune diseases as a possible factor that can evoke autoimmunity (33, 34).

**Figure 5.**
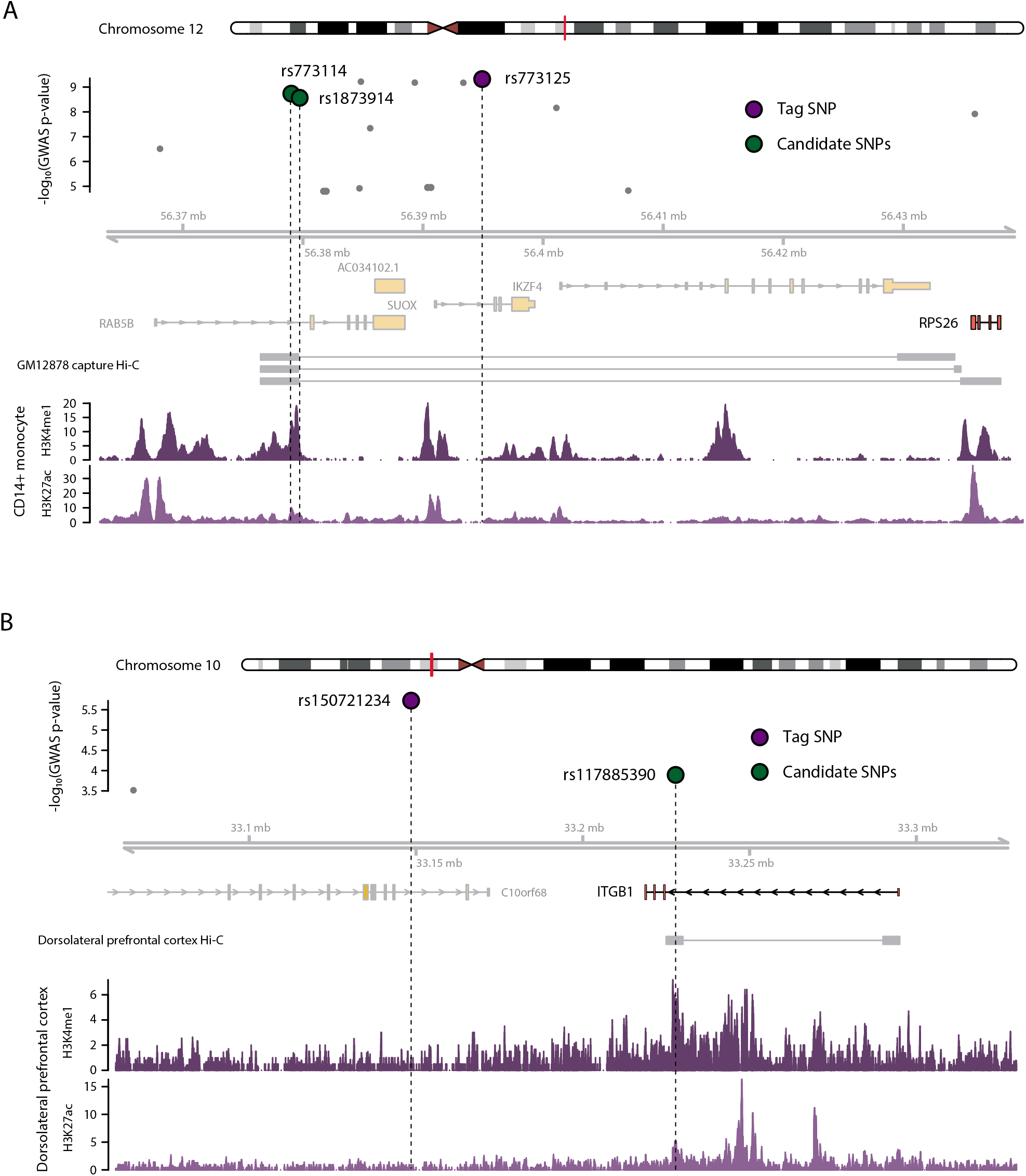
Examples of novel candidate causal SNPs. **(a)** Example for autoimmune diseases. Known GWAS SNP rs773125, located on chromosome 12, has the strongest association with RA. However, this SNP was a negative call from our prediction and not located in active regulatory regions in any immune-related tissues. The two positive SNPs (rs773114 and rs1873914) were located in an active regulatory region of CD14+ monocyte and connected to RPS26 according to GM12878 capture Hi-C data (32). **(b)** Example for psychiatric disorders. rs150721234 has the strongest association strength with SCZ in the LD block located on chromosome 10. However, this SNP was a negative call from our prediction and not located in active regulatory regions in any brain-related tissues. Instead, there was a positive SNP (rs117885390) that showed a lower association strength with SCZ. Dorsolateral prefrontal cortex Hi-C data (35) located this SNP in an enhancer region of ITGB1. This site was an active regulatory region of the dorsolateral prefrontal cortex tissue.

We were able to find similar examples in psychiatric disorders. For example, rs150721234 has the strongest association strength with the SCZ phenotype in the LD block located on chromosome 10 (Figure 5b). This SNP resides in an intron region of C10orf68, which has no clear role in association with SCZ. Moreover, rs150721234 is not located in active regulatory regions marked by H3K4me1 or H3K27ac in any types of brain-related tissues. Our prediction was negative on rs150721234. Instead, there was a positive SNP (rs117885390) that showed a lower association strength with SCZ. Dorsolateral prefrontal cortex Hi-C data (35) located this SNP in an enhancer region of ITGB1 (Figure 5b). This site was an active regulatory region of the dorsolateral prefrontal cortex. There were several studies showing that ITGB1 gene has a possible role during the development of schizophrenia (36, 37). These cases show that our method may be able to pinpoint functional variants that could be the actual cause of diseases among all variants associated with tag SNPs. Furthermore, these examples illustrate that once candidate variants other than tag SNPs are identified, one may be able to specify novel target genes that may shed light on the pathophysiology of the relevant phenotypes.

## DISCUSSION

Our prediction model is different from the conventional architecture of CNNs intended for image processing. For biological reasons, we perform onedimensional convolution while using a vector instead of a matrix for kernels. Only one-dimensional convolution is applicable for our purpose because genetic information is encoded in linear DNA strands. This type of convolution has been applied for predicting the sequence motifs of DNA- and RNA-binding proteins (8). While binding motifs are consecutive nucleotides that can be represented as a positional matrix, the order of SNPs along the chromosome does not carry meaningful biological information. This is why only vector kernels were applicable for our purpose. The power of our method stems from incorporating feature data that comes with external annotation. These features were not learned from DNA sequences *ab initio* but were incorporated on the basis of domain knowledge, which probably contributed to achieving high performance with a relatively small number of convolution layers. In addition, biological annotation helped with the validation and interpretation of prediction results.

Statistical approaches for fine-mapping are not applicable for rare variants because of limited power. In contrast to statistical fine-scale mapping, our functional prediction method is applicable to rare variants for which statistical association is difficult to estimate. This is important because the combined effects of rare variants may explain a significant proportion of genetic susceptibility to common diseases or traits (38–40). Expression quantitative loci with large effects detected in a human family were enriched with rare regulatory variants (41). A burden test for enrichment revealed a significant excess of rare regulatory variants at both extremes of gene expression, implicating their potential role in contributing to disease by driving high or low transcription (42). Our method can contribute to the identification and prioritization of rare variants.

## ACKNOWLEDGEMENTS

None.

