## Supplementary Figures and Information for "Convolutional neural network model to predict causal risk factors that share complex regulatory features"

**Supplementary Figure 1. Illustration of feature map construction.**

This figure illustrates how epigenomic data were processed into feature maps. When a SNP in an association block falls in a peak of epigenomic signal, the corresponding entry of the feature map is assigned 1 (colored cells). For example, SNP #3 in association block #1 overlaps with feature #1, #2, and #4, and is assigned 1 for these features in the feature map of this block. Features that did not map to any SNPs in >95% of association blocks (e.g., feature #3) were filtered out.

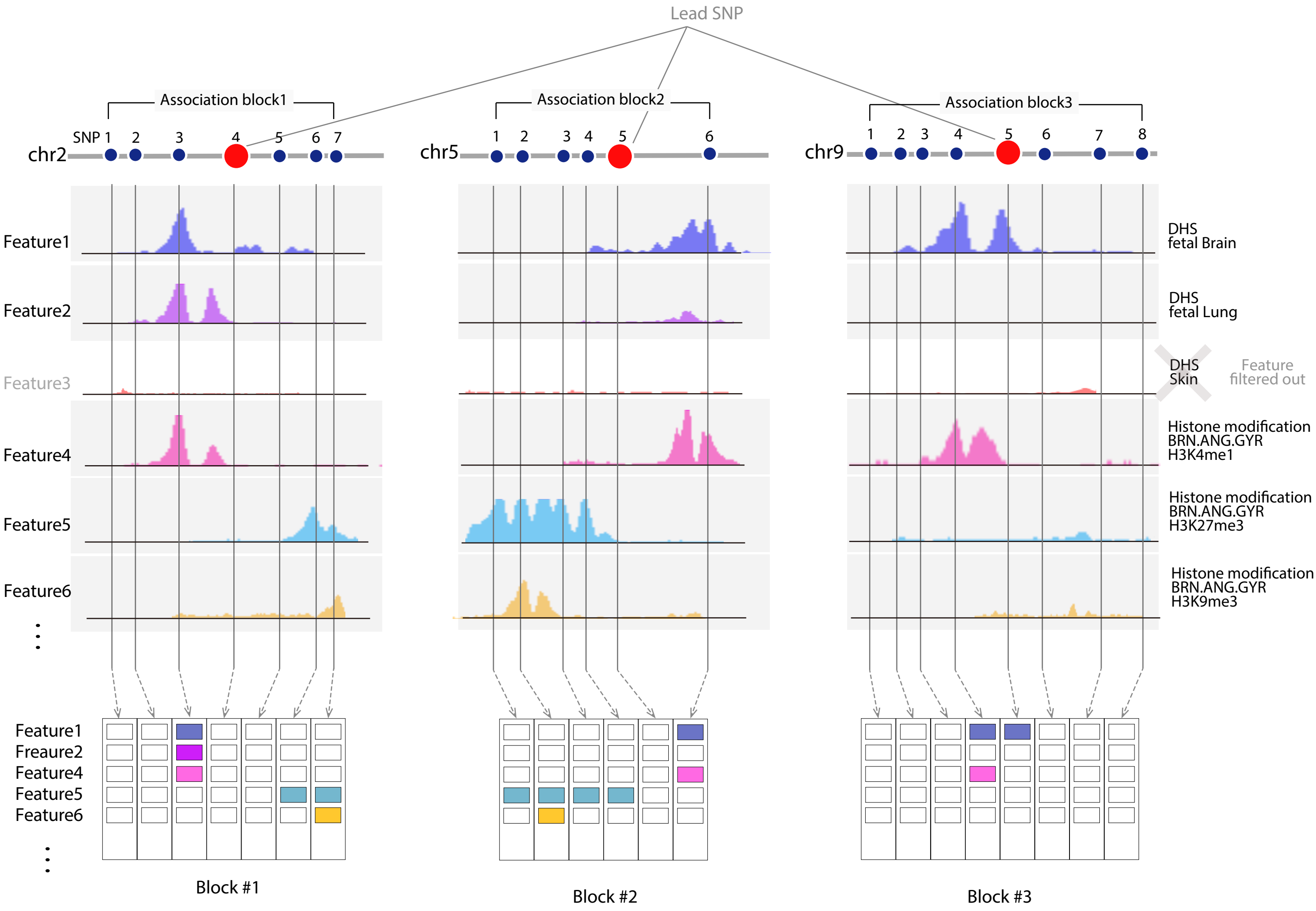

Lead SNP
Association block1
Association block2
Association block3
chr2
SNP 1 2 3 4 5 6 7
chr5
1 2 3 4 5 6
chr9
1 2 3 4 5 6 7 8
Feature1
Feature2
Feature3
Feature4
Feature5
Feature6
DHS fetal Brain
DHS fetal Lung
DHS Skin
Feature filtered out
Histone modification BRN.ANG.GYR H3K4me1
Histone modification BRN.ANG.GYR H3K27me3
Histone modification BRN.ANG.GYR H3K9me3
Feature1
Freaure2
Feature4
Feature5
Feature6
Block #1
Block #2
Block #3

**Supplementary Figure 2. Schematic of the learning processes.**

Association blocks in different chromosomes were used as the training, validation, and testing sets, separately. Out of 188,160 possible combinations, randomly chosen 500 hyperparameter sets were used. The best model was chosen based on the F1 score obtained from the validation set. Final performance was assessed on the testing set in terms of F1 and AUC.

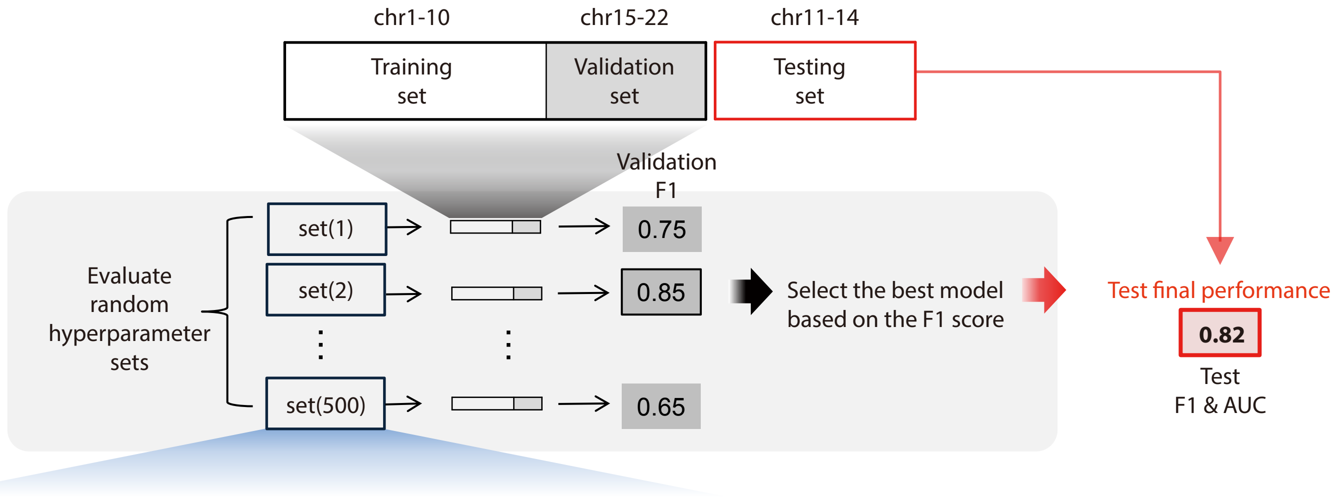


| Hyperparametsers | Search space |
|---|---|
| pretrain learning rate | [1.5, 1., 5e-01, 1e-01, 5e-02, 1e-02, 5e-03, 1e-03] |
| corruption level ( $\alpha$ ) | [0, 0.1, 0.2, 0.3, ..., 0.9] |
| learning rate( $\varepsilon$ ) | [1.5, 1., 5e-01, 1e-01, 5e-02, 1e-02, 5e-03, 1e-03] |
| $\lambda$ 1 | [0, 1e-05, 1e-04, 1e-03, 1e-02, 1e-01, 1.] |
| $\lambda$ 2 | [0, 1e-05, 1e-04, 1e-03, 1e-02, 1e-01, 1.] |
| Momentum ( $\mu$ ) | [0, 0.1, 0.2, ..., 0.5] |
| number of kernels ( $K$ ) | [50] |
| Batch size ( $B$ ) | [100] |

188,160 combinations in total

**Supplementary Figure 3. Validation of prediction results in terms of statistical association.** **(a)** Fraction of positive calls among the SNPs that were associated with each disease phenotype at $P < 0.001$ in comparison to that among the SNPs that were not associated ($P > 0.1$). **(b)** The average prediction score of the SNPs that were associated with each disease at $P < 0.001$ in comparison to that among the SNPs that were not associated ($P > 0.1$). To obtain the statistical significance of the mean differences, we sampled the same number of random SNPs as the associated SNPs, and repeated the sampling 10,000 times. **(a-b)** Because the learning was performed on associated SNPs in the training and validation sets, the correlation of association strength and prediction score is expected for these SNPs. Therefore, we tested the correlation by using SNPs in the testing set.

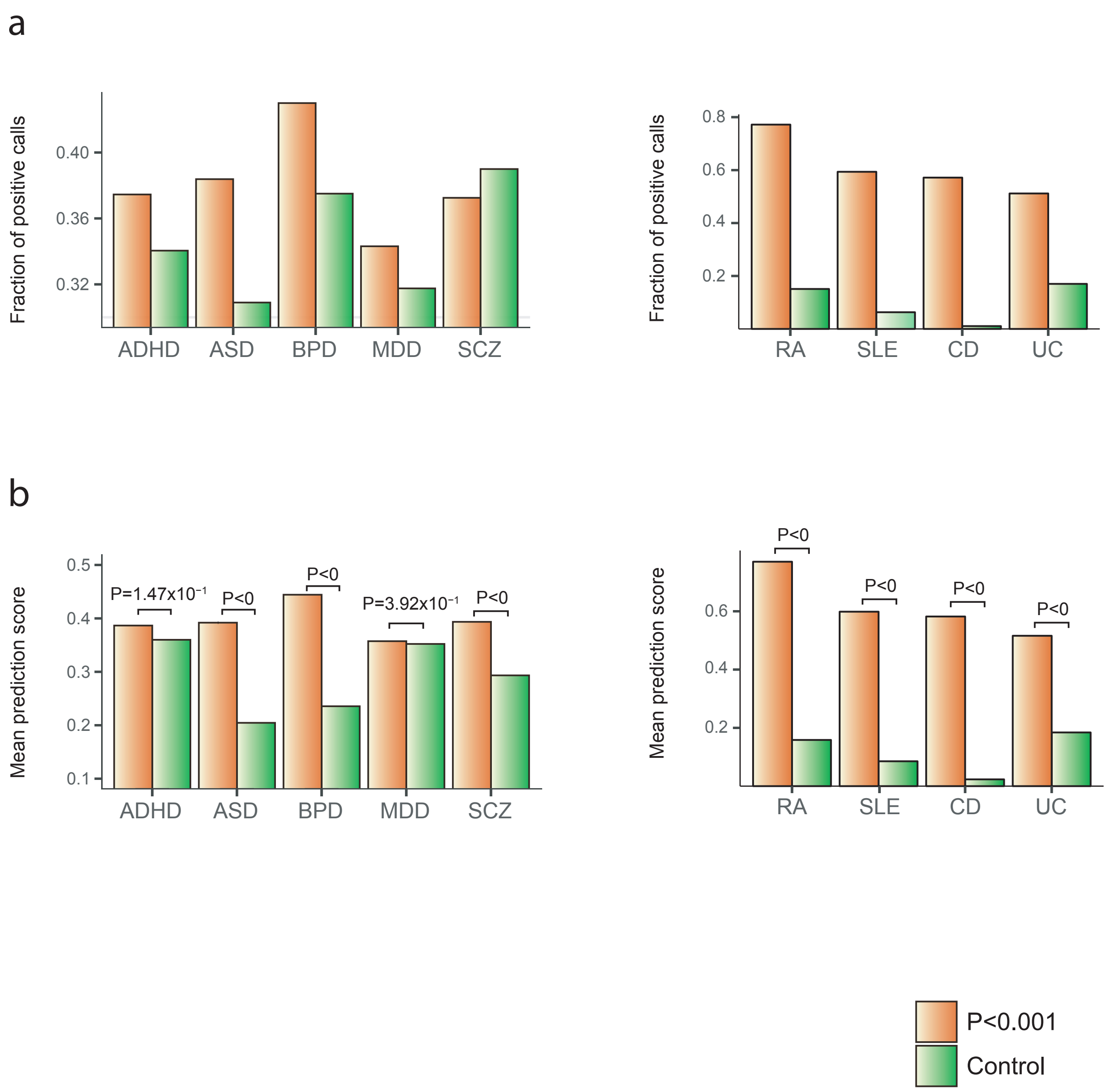

a
Fraction of positive calls
0.40
0.36
0.32
ADHD
ASD
BPD
MDD
SCZ
Fraction of positive calls
0.8
0.6
0.4
0.2
RA
SLE
CD
UC
b
Mean prediction score
0.5
0.4
0.3
0.2
0.1
P=1.47x$10^{-1}$
P<0
P<0
P=3.92x$10^{-1}$
P<0
ADHD
ASD
BPD
MDD
SCZ
Mean prediction score
0.6
0.4
0.2
P<0
P<0
P<0
P<0
RA
SLE
CD
UC
P<0.001
Control

**Supplementary Figure 4. Examples of prediction results in comparison with imputed statistical association.**

Each SNP in a chromosomal block is assigned a prediction score (red diamond on the left axis) and the P value of phenotypic association (blue dot on the right axis). These examples display cases in which the greatest prediction score was assigned to variants with strongest association. The chromosomal position of the lead SNP marking each block is shown on the upper left corner.

RA

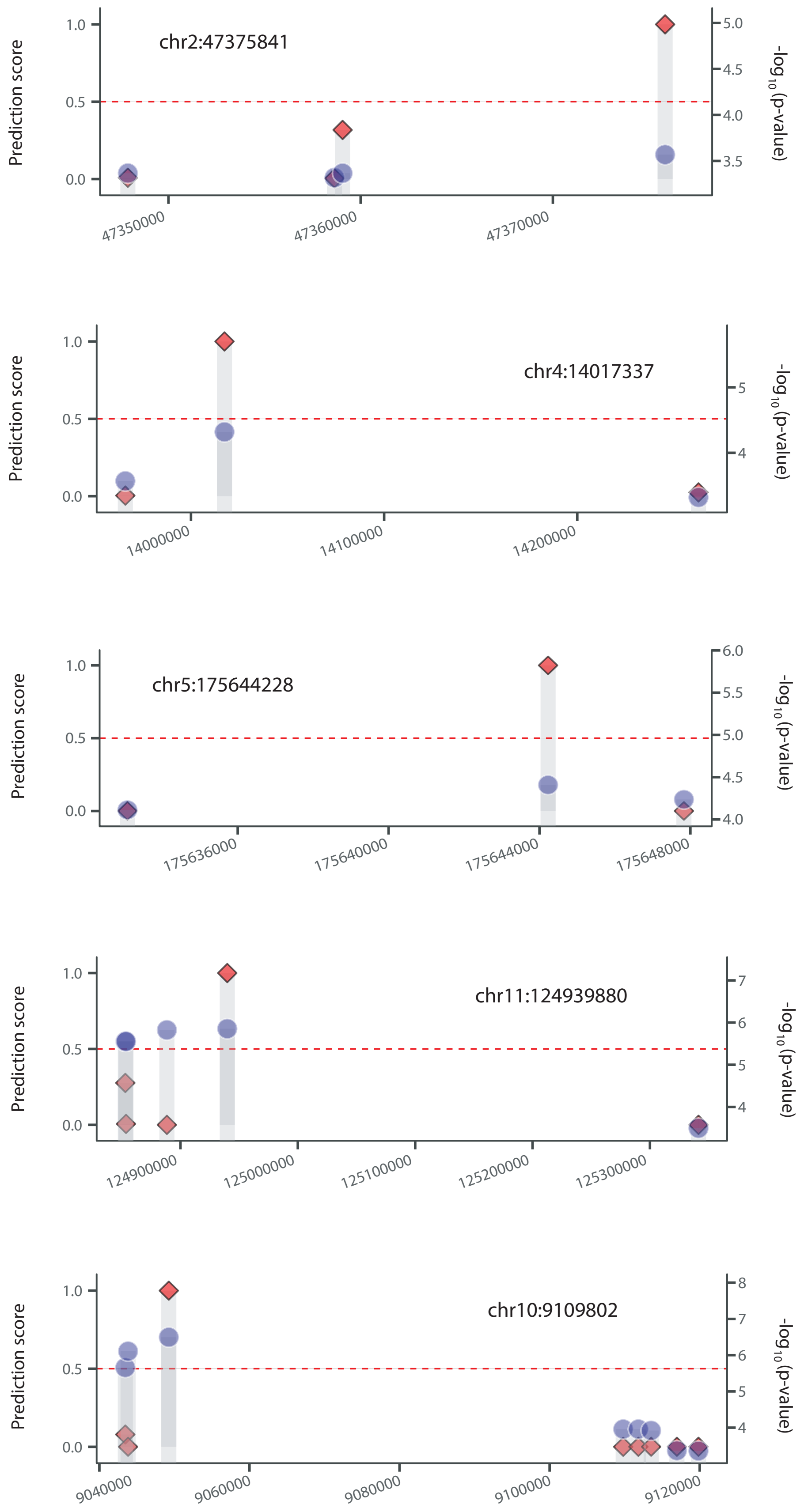
chr2:47375841
Prediction score
-log10(p-value)
1.0
0.5
0.0
5.0
4.5
4.0
3.5
47350000
47360000
47370000
chr4:14017337
Prediction score
-log10(p-value)
1.0
0.5
0.0
5
4
14000000
14100000
14200000
chr5:175644228
Prediction score
-log10(p-value)
1.0
0.5
0.0
6.0
5.5
5.0
4.5
4.0
175636000
175640000
175644000
175648000
chr11:124939880
Prediction score
-log10(p-value)
1.0
0.5
0.0
7
6
5
4
124900000
125000000
125100000
125200000
125300000
chr10:9109802
Prediction score
-log10(p-value)
1.0
0.5
0.0
8
7
6
5
4
9040000
9060000
9080000
9100000
9120000

SLE

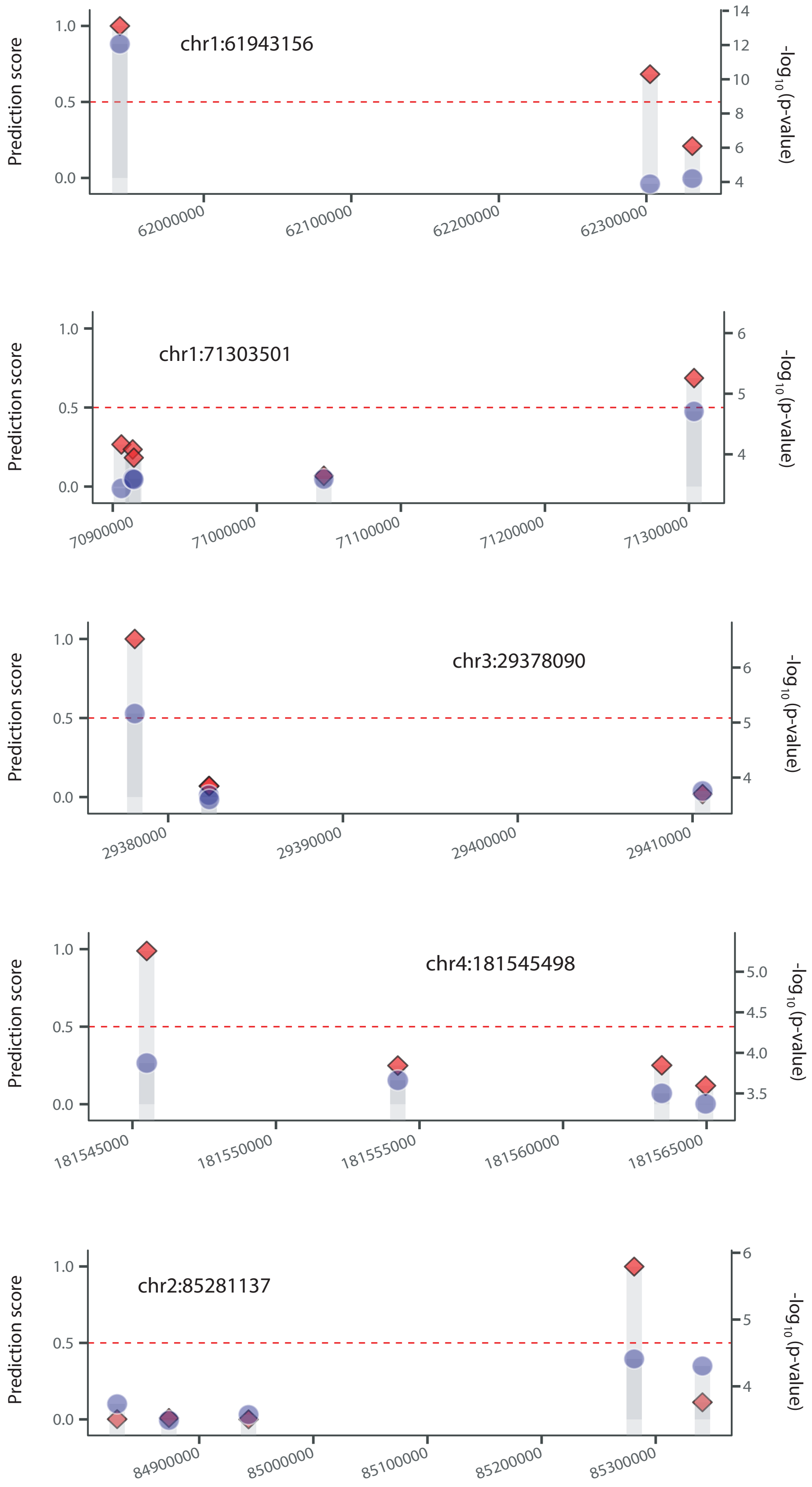
Prediction score
-log10(p-value)
chr1:61943156
62000000
62100000
62200000
62300000
chr1:71303501
70900000
71000000
71100000
71200000
71300000
chr3:29378090
29380000
29390000
29400000
29410000
chr4:181545498
181545000
181550000
181555000
181560000
181565000
chr2:85281137
84900000
85000000
85100000
85200000
85300000

CD

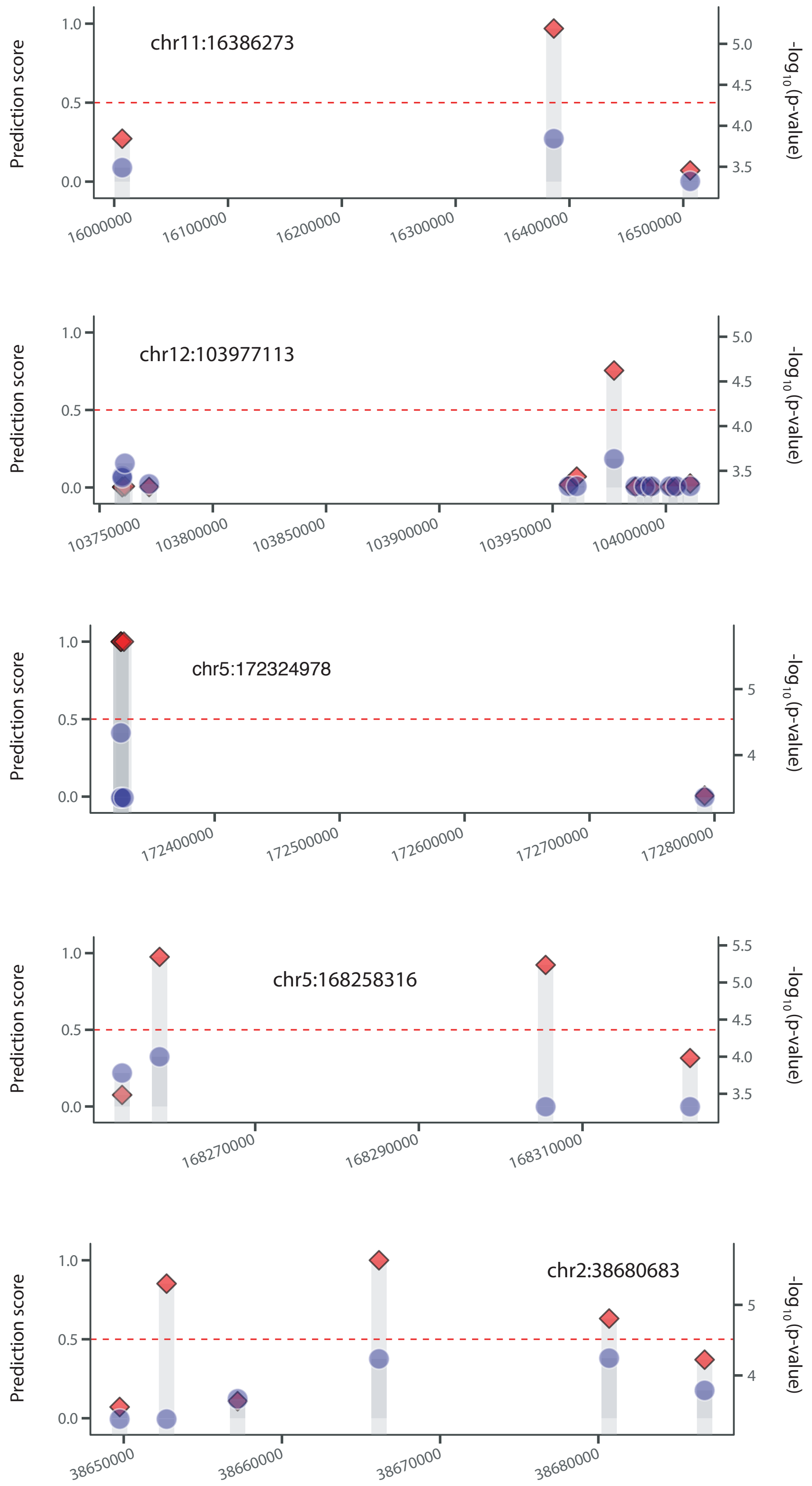
chr11:16386273
Prediction score
-log10(p-value)
16000000
16100000
16200000
16300000
16400000
16500000
chr12:103977113
103750000
103800000
103850000
103900000
103950000
104000000
chr5:172324978
172400000
172500000
172600000
172700000
172800000
chr5:168258316
168270000
168290000
168310000
chr2:38680683
38650000
38660000
38670000
38680000

UC

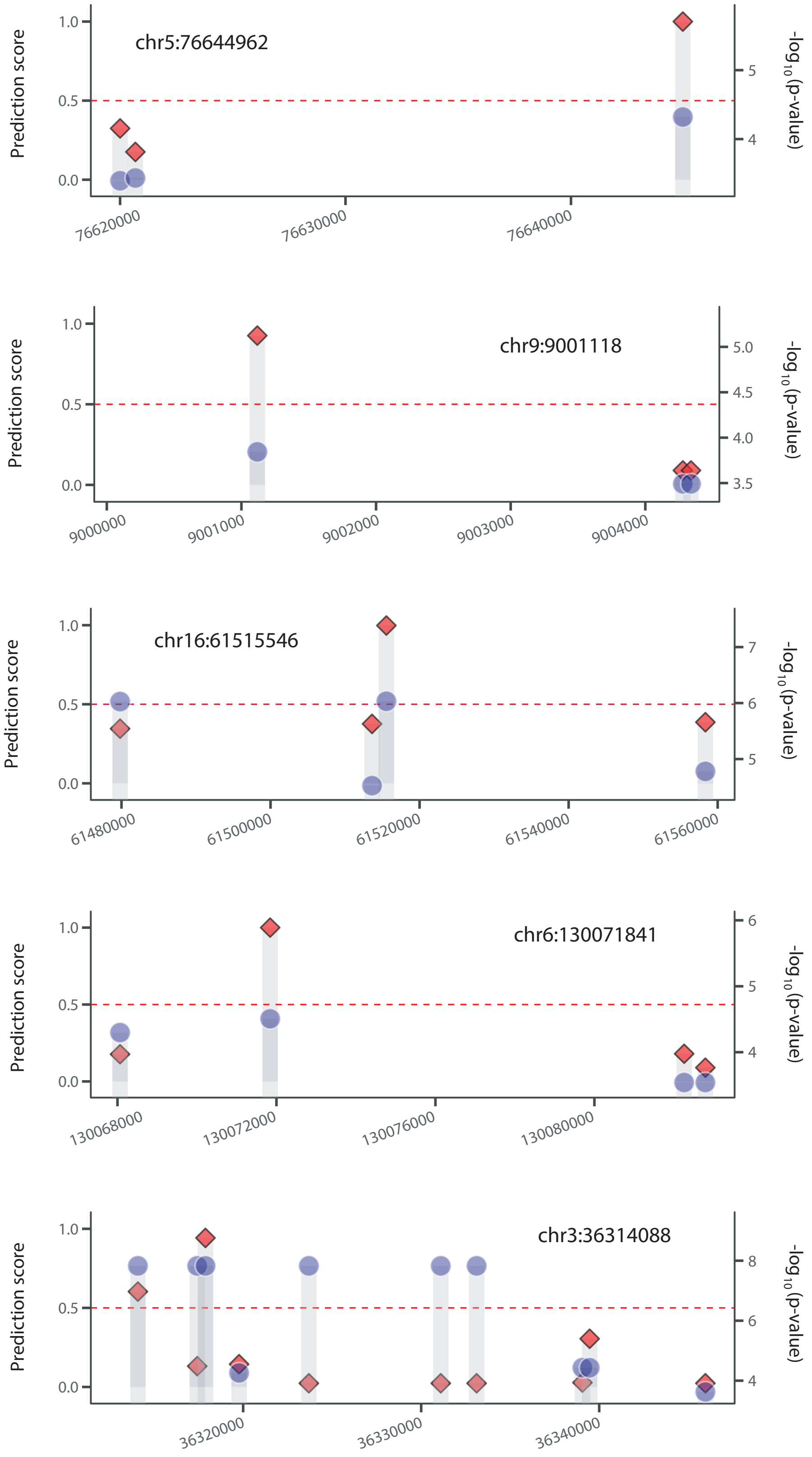
chr5:76644962
chr9:9001118
chr16:61515546
chr6:130071841
chr3:36314088
Prediction score
-log10(p-value)
1.0
0.5
0.0
76620000
76630000
76640000
9000000
9001000
9002000
9003000
9004000
61480000
61500000
61520000
61540000
61560000
130068000
130072000
130076000
130080000
36320000
36330000
36340000

ADHD

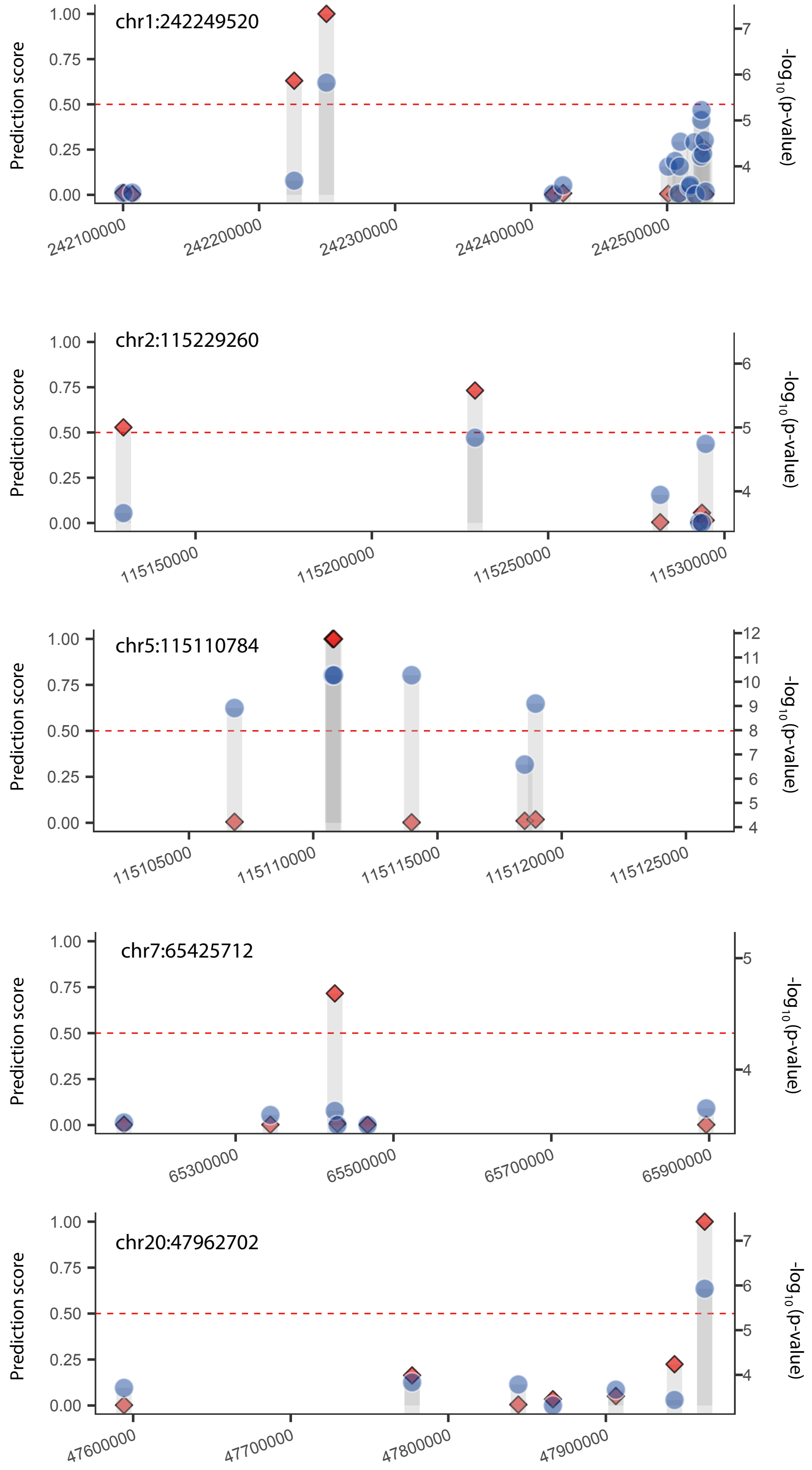
chr1:242249520
chr2:115229260
chr5:115110784
chr7:65425712
chr20:47962702
Prediction score
-log10(p-value)
1.00
0.75
0.50
0.25
0.00
242100000
242200000
242300000
242400000
242500000
115150000
115200000
115250000
115300000
115105000
115110000
115115000
115120000
115125000
65300000
65500000
65700000
65900000
47600000
47700000
47800000
47900000

ASD

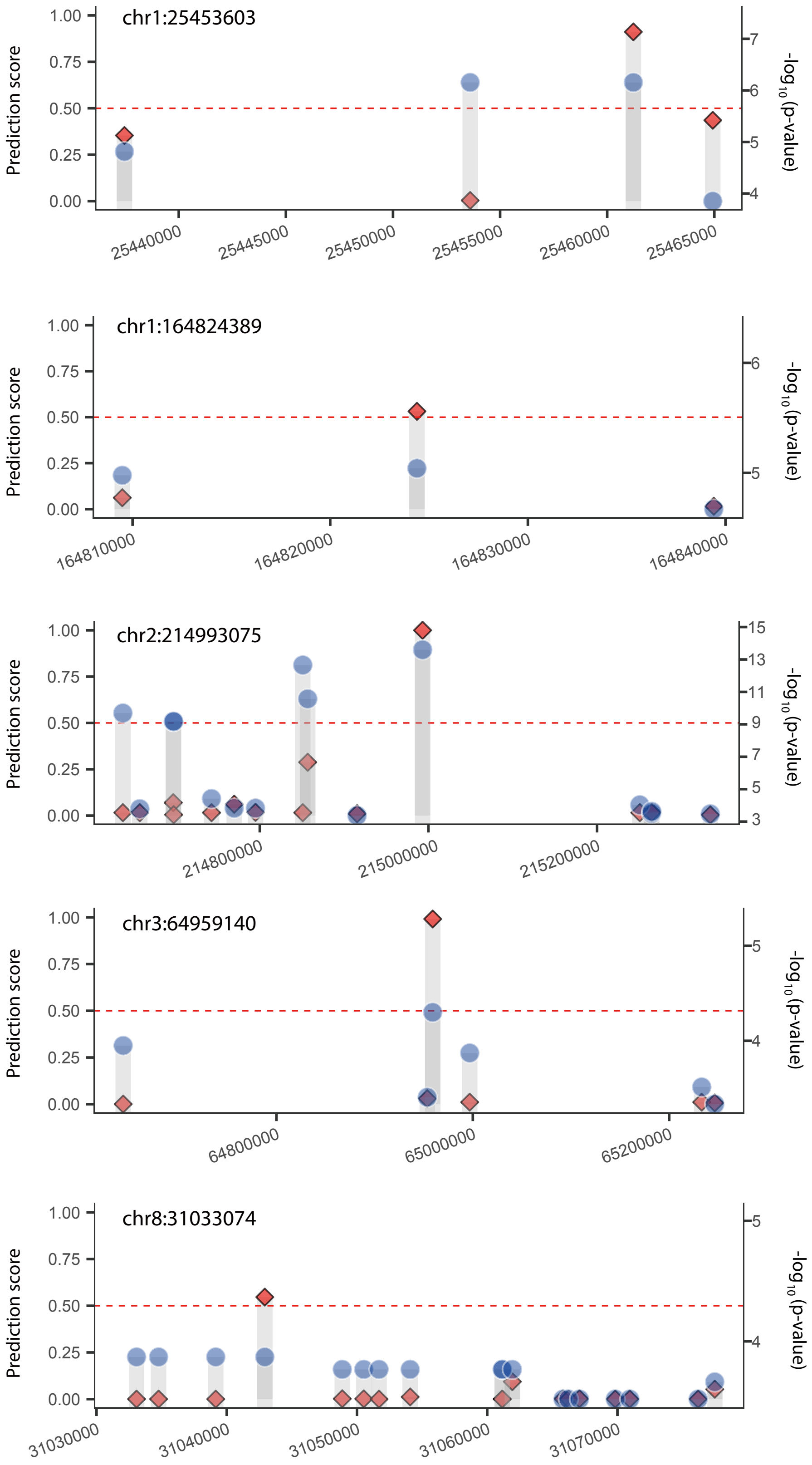
chr1:25453603
Prediction score
1.00
0.75
0.50
0.25
0.00
25440000
25445000
25450000
25455000
25460000
25465000
7
6
5
4
-log10(p-value)
chr1:164824389
Prediction score
1.00
0.75
0.50
0.25
0.00
164810000
164820000
164830000
164840000
6
5
-log10(p-value)
chr2:214993075
Prediction score
1.00
0.75
0.50
0.25
0.00
214800000
215000000
215200000
15
13
11
9
7
5
3
-log10(p-value)
chr3:64959140
Prediction score
1.00
0.75
0.50
0.25
0.00
64800000
65000000
65200000
5
4
-log10(p-value)
chr8:31033074
Prediction score
1.00
0.75
0.50
0.25
0.00
31030000
31040000
31050000
31060000
31070000
5
4
-log10(p-value)

BPD

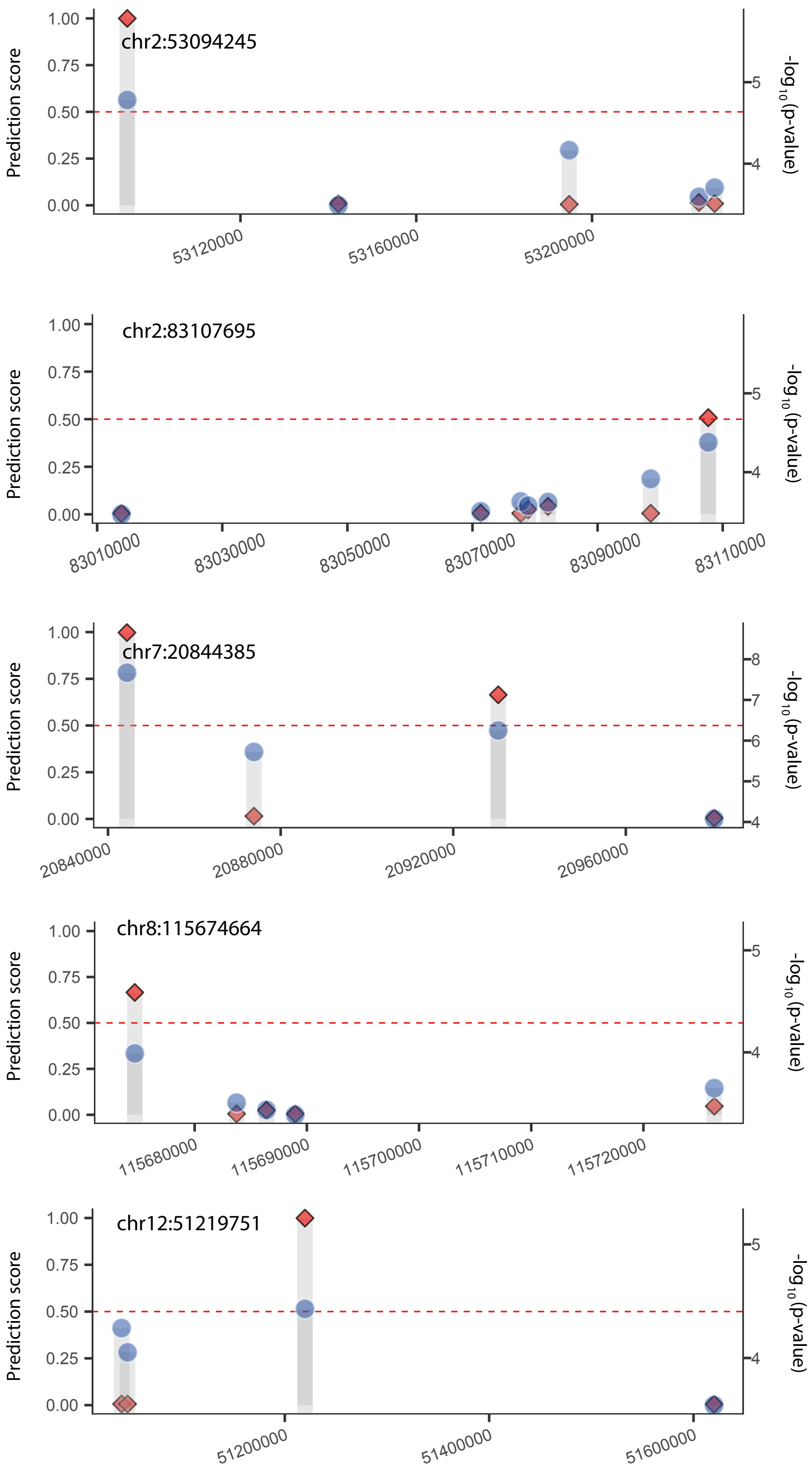

chr2:53094245
Prediction score
1.00
0.75
0.50
0.25
0.00
53120000
53160000
53200000
-log10(p-value)
5
4
chr2:83107695
83010000
83030000
83050000
83070000
83090000
83110000
chr7:20844385
20840000
20880000
20920000
20960000
8
7
6
5
4
chr8:115674664
115680000
115690000
115700000
115710000
115720000
chr12:51219751
51200000
51400000
51600000

MDD

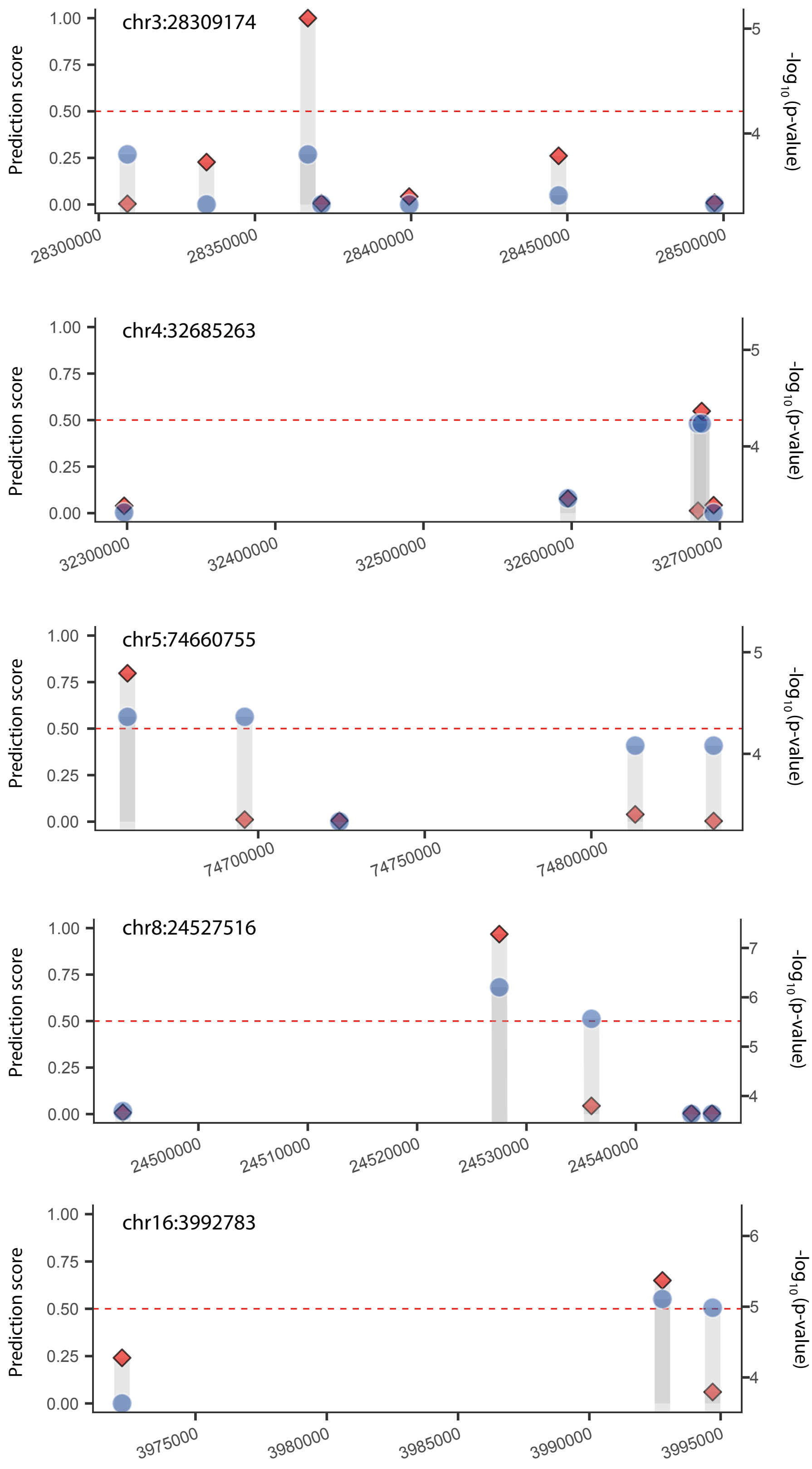
chr3:28309174
Prediction score
1.00
0.75
0.50
0.25
0.00
28300000
28350000
28400000
28450000
28500000
5
4
-log10(p-value)
chr4:32685263
32300000
32400000
32500000
32600000
32700000
chr5:74660755
74700000
74750000
74800000
chr8:24527516
24500000
24510000
24520000
24530000
24540000
7
6
chr16:3992783
3975000
3980000
3985000
3990000
3995000

SCZ

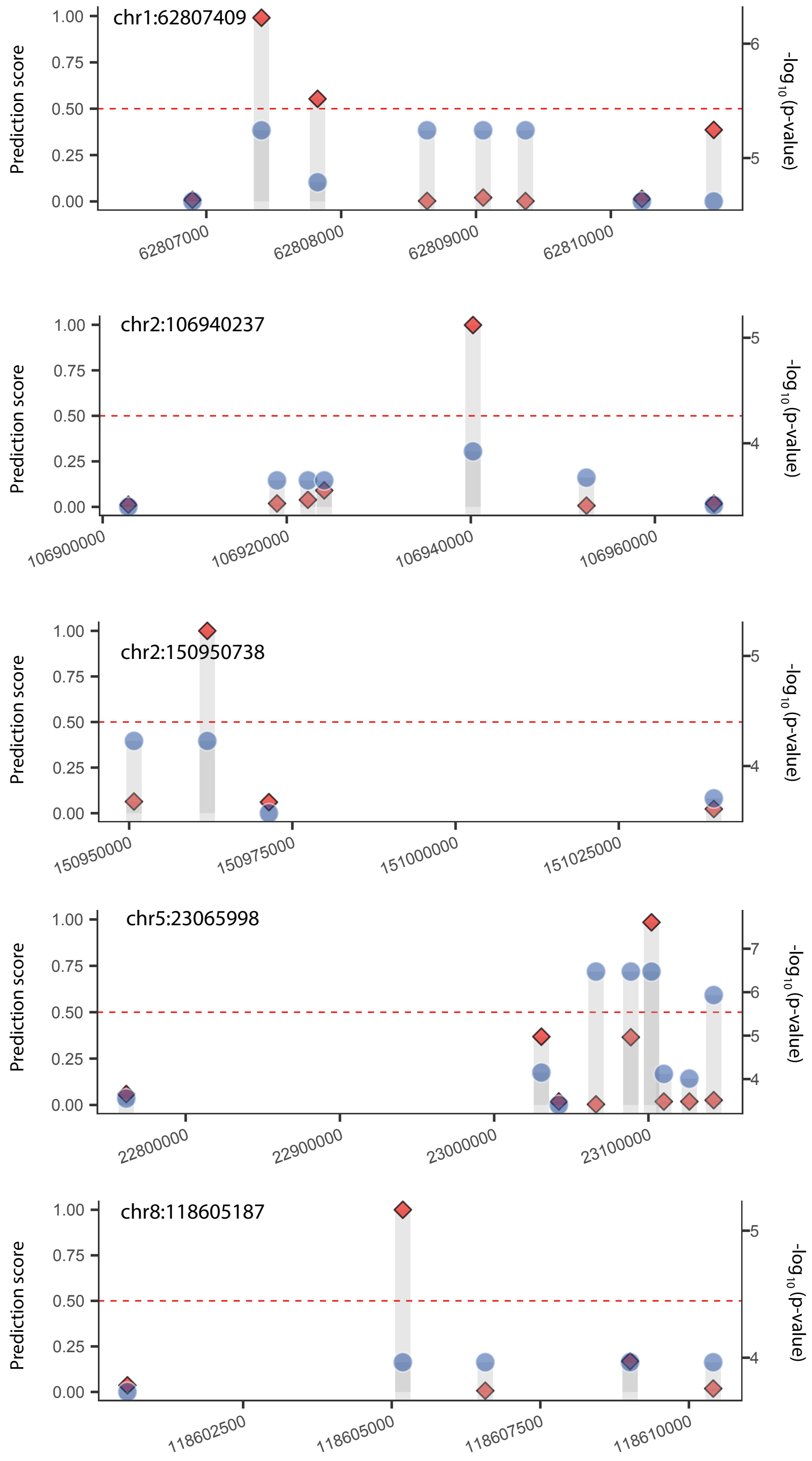
chr1:62807409
chr2:106940237
chr2:150950738
chr5:23065998
chr8:118605187
Prediction score
-log10(p-value)
1.00
0.75
0.50
0.25
0.00
62807000
62808000
62809000
62810000
106900000
106920000
106940000
106960000
150950000
150975000
151000000
151025000
22800000
22900000
23000000
23100000
118602500
118605000
118607500
118610000

**Supplementary Figure 5. Functional enrichment of target genes of positive SNPs.**

For target gene mapping, we used the dataset for distal enhancer-to-promoter connections based on the correlation of the sequencing tag density between distal DHSs and proximal DHSs across cell types for psychiatric disorders, and multiple long-range interaction data consist of capture Hi-C, ChIA-PET, and IM-PET for autoimmune diseases. Connections with Pearson correlation coefficient > 0.7 were used. Gene set enrichment analysis was run for the identified genes by using Enrichr. Shown here are the most significant terms related to the given psychiatric disorder or general neural function. The numbers attached to each term indicate the database resource from which the term has been derived:

1) dbGaP
2) KEGG pathway
3) Gene Ontology
4) Reactome
5) Disease perturbations from GEO

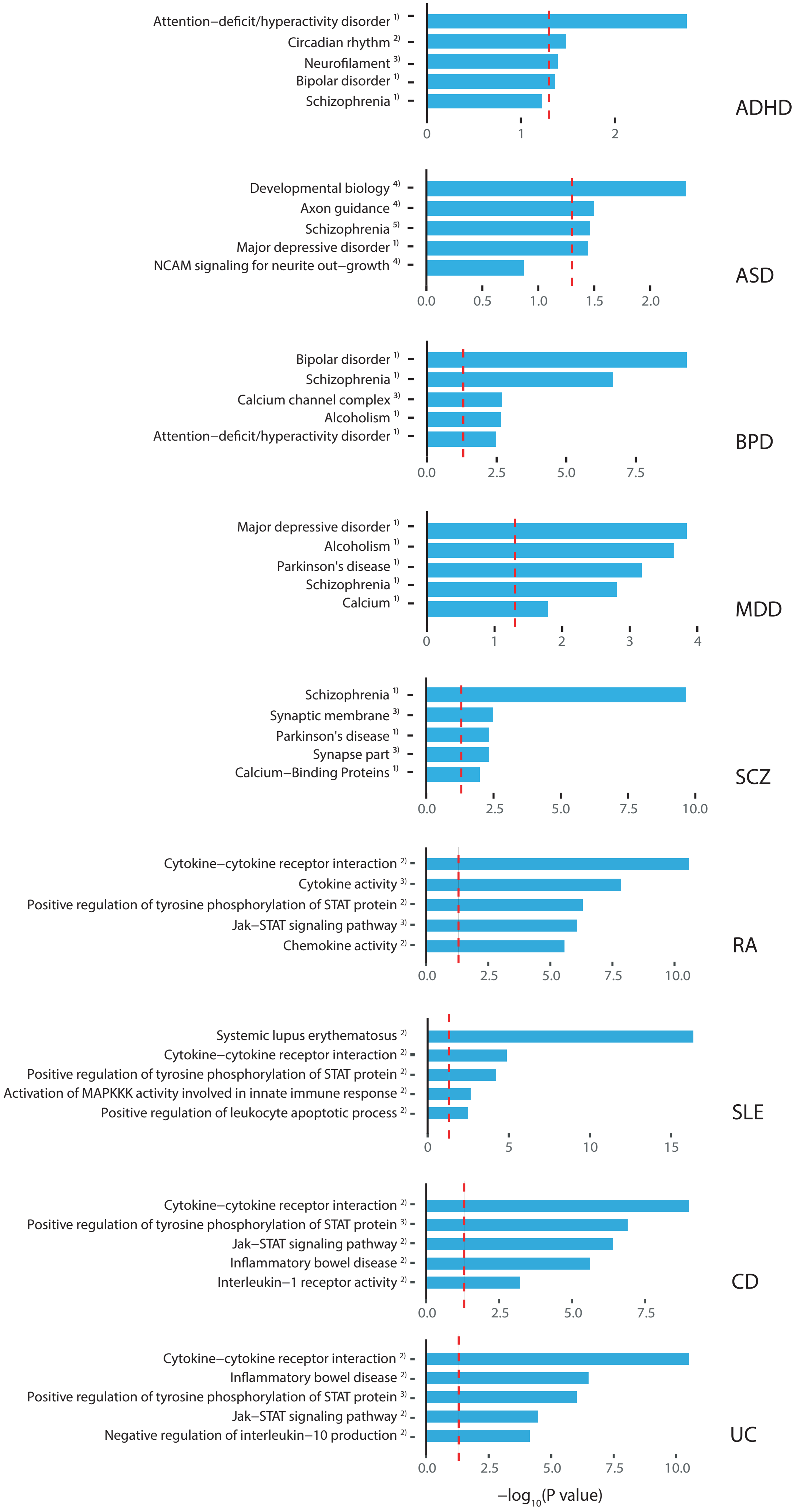

Attention–deficit/hyperactivity disorder 1)
Circadian rhythm 2)
Neurofilament 3)
Bipolar disorder 1)
Schizophrenia 1)
0
1
2
ADHD
Developmental biology 4)
Axon guidance 4)
Schizophrenia 5)
Major depressive disorder 1)
NCAM signaling for neurite out–growth 4)
0.0
0.5
1.0
1.5
2.0
ASD
Bipolar disorder 1)
Schizophrenia 1)
Calcium channel complex 3)
Alcoholism 1)
Attention–deficit/hyperactivity disorder 1)
0.0
2.5
5.0
7.5
BPD
Major depressive disorder 1)
Alcoholism 1)
Parkinson's disease 1)
Schizophrenia 1)
Calcium 1)
0
1
2
3
4
MDD
Schizophrenia 1)
Synaptic membrane 3)
Parkinson's disease 1)
Synapse part 3)
Calcium–Binding Proteins 1)
0.0
2.5
5.0
7.5
10.0
SCZ
Cytokine–cytokine receptor interaction 2)
Cytokine activity 3)
Positive regulation of tyrosine phosphorylation of STAT protein 2)
Jak–STAT signaling pathway 3)
Chemokine activity 2)
0.0
2.5
5.0
7.5
10.0
RA
Systemic lupus erythematosus 2)
Cytokine–cytokine receptor interaction 2)
Positive regulation of tyrosine phosphorylation of STAT protein 2)
Activation of MAPKKK activity involved in innate immune response 2)
Positive regulation of leukocyte apoptotic process 2)
0
5
10
15
SLE
Cytokine–cytokine receptor interaction 2)
Positive regulation of tyrosine phosphorylation of STAT protein 3)
Jak–STAT signaling pathway 2)
Inflammatory bowel disease 2)
Interleukin–1 receptor activity 2)
0.0
2.5
5.0
7.5
CD
Cytokine–cytokine receptor interaction 2)
Inflammatory bowel disease 2)
Positive regulation of tyrosine phosphorylation of STAT protein 3)
Jak–STAT signaling pathway 2)
Negative regulation of interleukin–10 production 2)
0.0
2.5
5.0
7.5
10.0
UC
–log10(P value)

**Supplementary Figure 6.** Enrichment of positive calls for autoimmune diseases for the BLUEPRINT epigenome data for various immune cell types. Positive calls were more enriched in the regulatory regions of lymphocyte lineages rather than granulocytes. Shown here are regulatory regions marked by H3K27ac .

H3K27ac

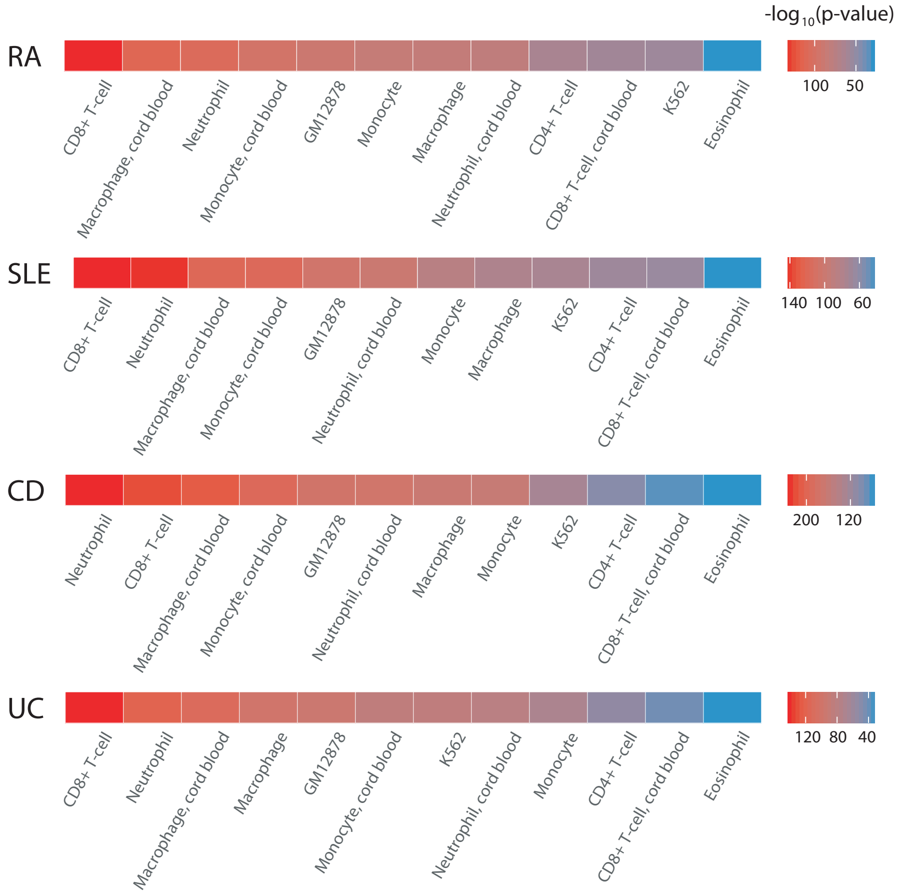

-log10(p-value)
RA
CD8+ T-cell
Macrophage, cord blood
Neutrophil
Monocyte, cord blood
GM12878
Monocyte
Macrophage
Neutrophil, cord blood
CD4+ T-cell
CD8+ T-cell, cord blood
K562
Eosinophil
100 50
SLE
CD8+ T-cell
Neutrophil
Macrophage, cord blood
Monocyte, cord blood
GM12878
Neutrophil, cord blood
Monocyte
Macrophage
K562
CD4+ T-cell
CD8+ T-cell, cord blood
Eosinophil
140 100 60
CD
Neutrophil
CD8+ T-cell
Macrophage, cord blood
Monocyte, cord blood
GM12878
Neutrophil, cord blood
Macrophage
Monocyte
K562
CD4+ T-cell
CD8+ T-cell, cord blood
Eosinophil
200 120
UC
CD8+ T-cell
Neutrophil
Macrophage, cord blood
Macrophage
GM12878
Monocyte, cord blood
K562
Neutrophil, cord blood
Monocyte
CD4+ T-cell
CD8+ T-cell, cord blood
Eosinophil
120 80 40

**Supplementary Figure 7. Regulatory effects of positive SNPs.**

**(a)** Fraction of variants causing allelic imbalance in chromatin accessibility at TF binding sites. The degree of allelic imbalance was measured based on the number of DHS footprint reads carrying each allele. We tested whether the ratio of reads from each allele (allele 1:allele 2) deviated from 50:50 by using the binomial test and multiple testing adjustment. We then compared the fraction of significantly imbalanced variants between positive and negative calls. **(b)** Association of the genotypes with the expression level of putative target genes compared between positive and negative SNPs. Expression association was calculated for all genes in the association block of the given SNP by using RNA-seq data for 465 lymphoblastoid cells whose nucleotide-resolution genotypes are available from the 1000 Genomes Project. The gene with the strongest association level was regarded as the target gene.

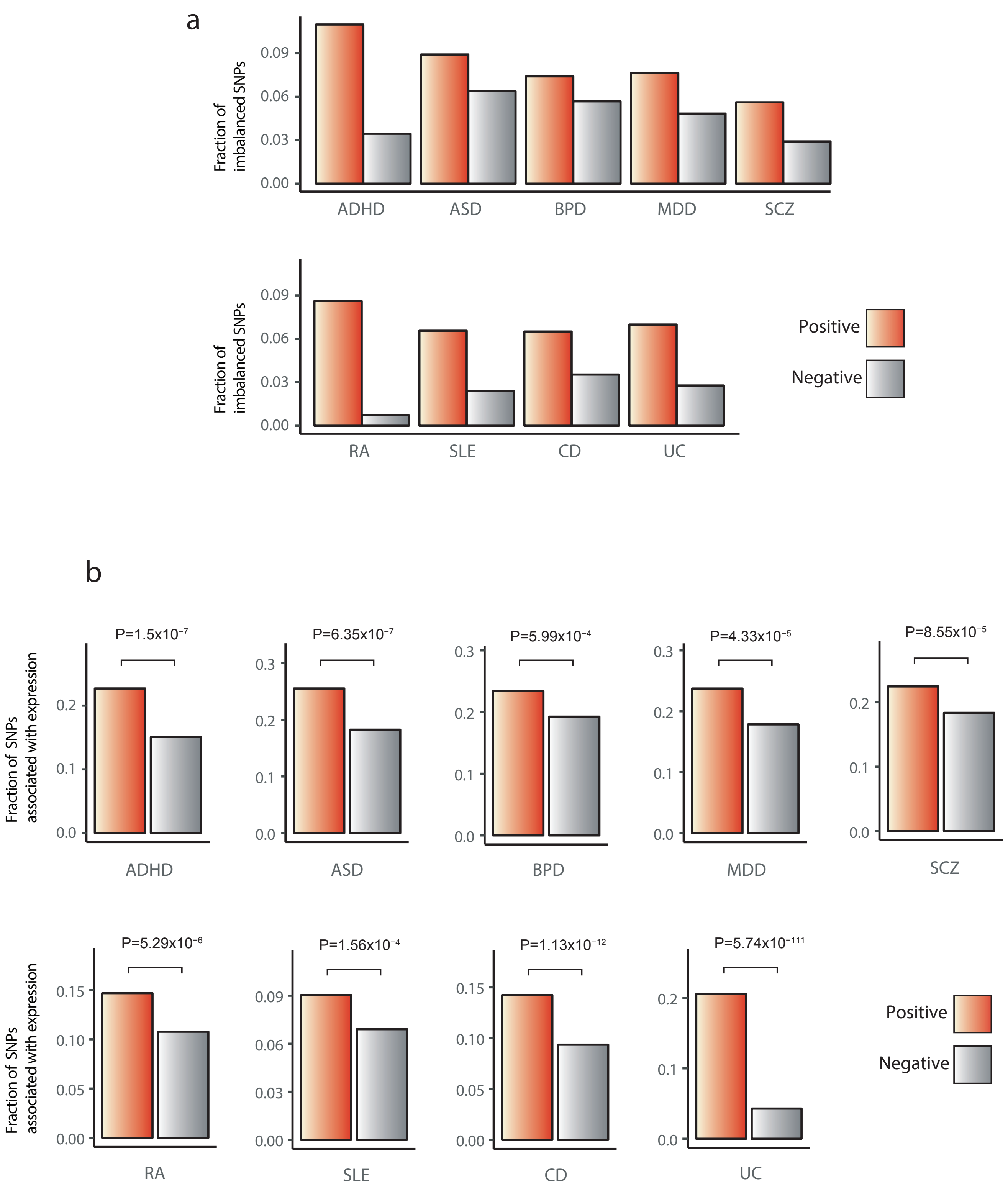
a
Fraction of imbalanced SNPs
0.09
0.06
0.03
0.00
ADHD
ASD
BPD
MDD
SCZ
RA
SLE
CD
UC
Positive
Negative
b
Fraction of SNPs associated with expression
P=1.5x10⁻⁷
P=6.35x10⁻⁷
P=5.99x10⁻⁴
P=4.33x10⁻⁵
P=8.55x10⁻⁵
P=5.29x10⁻⁶
P=1.56x10⁻⁴
P=1.13x10⁻¹²
P=5.74x10⁻¹¹¹
ADHD
ASD
BPD
MDD
SCZ
RA
SLE
CD
UC
Positive
Negative

**Supplementary Figure 8. Evolutionary conservation of genomic regions with positive call. (a)** The average PhastCons score for primates of regions overlapping positive (red) vs. negative (gray) prediction. **(b)** The ratio of the odds of positive SNPs in conserved regions to their odds in non-conserved regions. For the conserved regions, we searched association blocks for the mammal or vertebrate PhastCons score > 0.5. The odds of positive SNPs were computed as the ratio of the positive to negative SNPs in the conserved or non-conserved regions. Shown is the odds ratio together with its 95% confidence interval and P value.

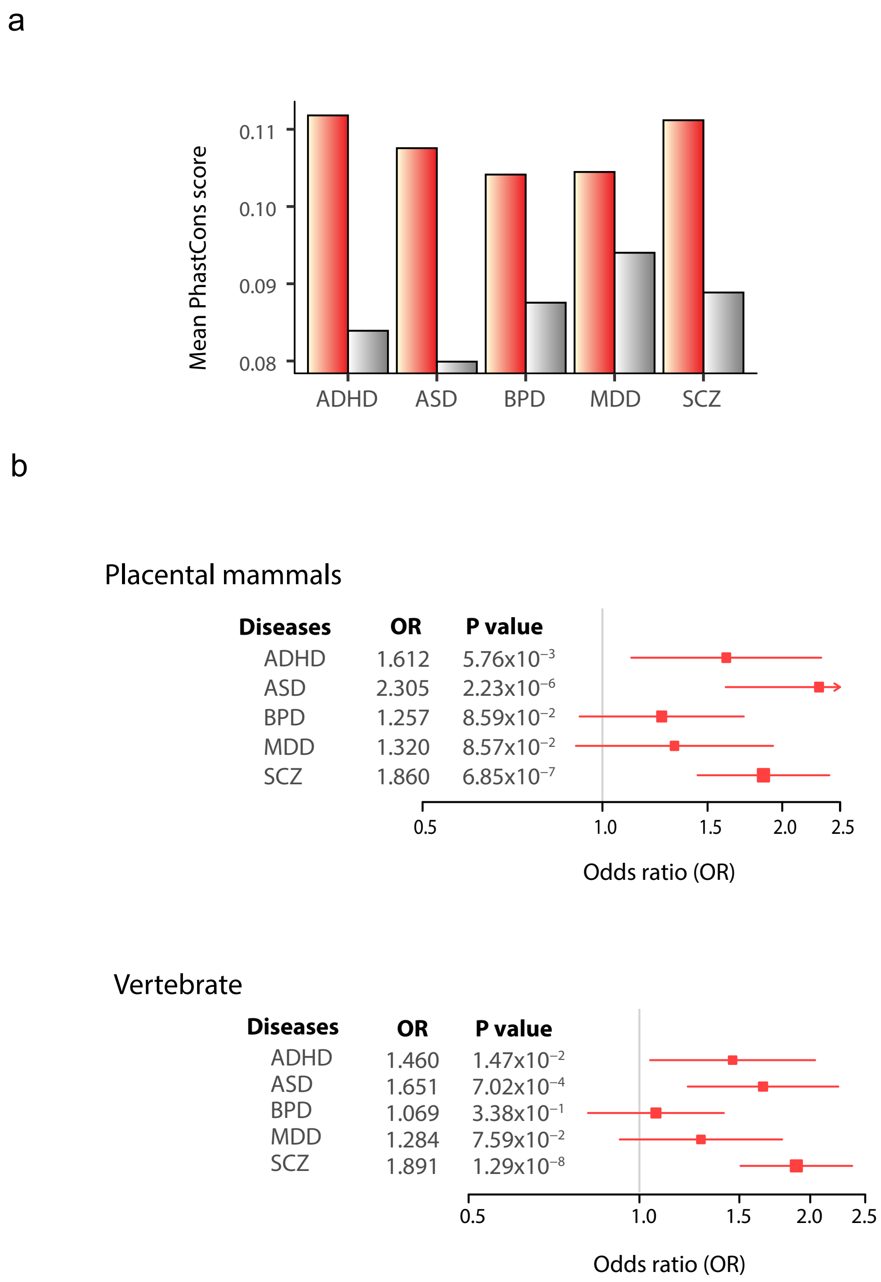
a
Mean PhastCons score
0.11
0.10
0.09
0.08
ADHD
ASD
BPD
MDD
SCZ
b
Placental mammals
Diseases
OR
P value
ADHD 1.612 5.76x10−3
ASD 2.305 2.23x10−6
BPD 1.257 8.59x10−2
MDD 1.320 8.57x10−2
SCZ 1.860 6.85x10−7
0.5
1.0
1.5
2.0
2.5
Odds ratio (OR)
Vertebrate
Diseases
OR
P value
ADHD 1.460 1.47x10−2
ASD 1.651 7.02x10−4
BPD 1.069 3.38x10−1
MDD 1.284 7.59x10−2
SCZ 1.891 1.29x10−8
0.5
1.0
1.5
2.0
2.5
Odds ratio (OR)

### Supplementary Information

#### Supplementary Methods

- Supplementary Methods S1: CNN model training processes
- Supplementary Methods S2: Functional analyses of predicted variants
- Supplementary References

#### Supplementary Methods S1: CNN model training proceses

#### Model training

##### - Training, validation, testing

To avoid overlapping of input features between the training, validation, and testing sets, we split our inputs by chromosomes. Chromosomes 1-10 were used as the training set, and chromosomes 11-14 were used as the testing set to report final performance levels. Chromosomes 15-22 were used as the validation set to select the best hyperparameters. Considering the length of chromosomes, approximately 3/5, 1/5, and 1/5 of the input data were assigned as the training, validation, and testing sets, respectively. The training set was broken into two subsets. One subset (chromosomes 1-7) was used for updating the model parameters and the other (chromosomes 8-10) was used for early stopping.

##### - Regularization

The training processes were aimed to minimize the LOSS function,

$$\mathrm{LOSS} = \mathrm{NLL} + \lambda_1 \|w\|_1 + \lambda_2 \|w\|_2,$$

where $\lambda_1 \|w\|_1 + \lambda_2 \|w\|_2$ represents the regularization term of the elastic net that is used to control overfitting. Specifically, $\|\cdot\|_p$ denotes the $L_p$ norm and defined as

$$\|w\|_p = (\sum_{j=0}^{|w|} |w_j|^p)^{\frac{1}{p}}.$$

$\lambda_p$ is a hyperparameter that controls the relative contribution of each regularization term. As implied in the above formula, the elastic net is a combination of lasso ( $L_1$ ) and ridge ( $L_2$ ) regularization on the weights of the model. The elastic net is known to outperform $L_1$ or $L_2$ regularization alone and to be especially useful when the number of predictors is large (1).

#### - Momentum

The momentum method is a technique to accelerate the gradient descent process by accumulating a velocity ( $v$ ) in the direction of stable and persistent reduction of the loss function (2). Given a loss function to be minimized, the momentum is given as

$$\theta_{t+1} = \theta_t + v_{t+1},$$

$$v_{t+1} = \mu v_t - \varepsilon \nabla(\mathrm{LOSS}(\theta^t)),$$

where $\varepsilon$ is the learning rate, $\mu$ is the momentum coefficient, and $\nabla(\mathrm{LOSS}(\theta^t))$ is the gradient at $\theta^t$. The value of $v_0$ was set to 0.

#### - Early stopping

As a measure to prevent overfitting, all models were trained only to the point when performance reached a plateau. We defined a variable called *patience* and initially set the value to 200. We then trained the model while keeping the number of training epochs below *patience*. Whenever performance improvement was observed, we updated *patience* to $\max(\text{initial } patience, \text{current epoch} \times 2)$ and saved the current model as the best. The threshold of improvement was defined as

$$F1 \text{ score of the best model} \times (1 + 0.002).$$

Because the numbers of the true and false cases are imbalanced, the *F1* score was thought to be more suitable for the evaluation of model performance than AUC (3). Therefore, we evaluated model performance based on the *F1* score, which is the harmonic mean of precision and recall:

$$F1 = \frac{2}{\frac{1}{\mathrm{Precision}} + \frac{1}{\mathrm{Recall}}},$$

$$\text{where } \mathrm{Precision} = \frac{\mathrm{TP}}{\mathrm{TP} + \mathrm{FP}}$$

$$\text{and} \quad \text{Recall} = \frac{\text{TP}}{\text{TP} + \text{FN}}.$$

TP, FP, and FN stand for true positives, false positives, and false negatives, respectively.

##### Pre-training by autoencoder

Autoencoder is an unsupervised learning component for deep learning that functions as a guide to the following fine-tuning process by assigning an optimal starting point where training data is better generalized (4–6). To allow the model to learn more robust features, the denoising autoencoder (DA) trains the model to reconstruct the input from a corrupted version of it. The DA was introduced to our model to pre-train the *K* filters. For pre-training by the DA, chromosomes 1-7 were used.

Our DA takes vectors of *M* features of each SNP as input:

$$x_m = \begin{cases} 1 & \text{if the } m\text{-}th \text{ feature is associated with the SNP} \\ & (\text{for all } 1 \le m \le M) \\ 0 & \text{otherwise} \end{cases}.$$

Here, each input vector ($x_m$) corresponds to each column of the input matrix $X_{mn}$ mentioned in the main text. The DA was trained to reconstruct $x$ from a stochastically corrupted version of $x$ ($\tilde{x}$). In other words, the DA learns an encoder $h(\cdot)$ and a decoder $g(\cdot)$ whose composition approaches $g(h(\tilde{x})) \approx x$ in the training set (4). We generated the corrupted version of input ($\tilde{x}$) by randomly masking entries with zero at every training step:

$$\tilde{x}_m = s_m x_m$$

$$\text{and} \quad s \sim \text{Bernoulli}(\alpha),$$

where $\alpha$ is the corruption level, and all mask variables ($s_m$) are sampled from Bernoulli distribution each time. The DA takes $\tilde{x}$ and maps it to a hidden representation $h(\tilde{x})$:

$$h(\tilde{x}) = \text{ReLU}(\tilde{x}W + b),$$

where $W$ is the weight matrix with a size of $M \times K$ and $b$ is the bias vector with a length of *K*. We set the number of the hidden nodes to *K*, which equals the number of kernels to be used for fine-tuning. The latent representation $h(\tilde{x})$ is then mapped back to $z$ given as

$$z = g(h(\tilde{x})),$$

$$z = \mathrm{sigmoid}(h(\tilde{x})W^T + b'),$$

where $b'$ is the bias vector with a length of *M*. The model parameters ($W$, $b$, and $b'$) were optimized to minimize the cross-entropy given as

$$C(x,z) = -\frac{1}{B}\sum_{s=1}^{B}(x^s \log z^s + (1-x^s)\log(1-z^s)),$$

where *B* is the mini-batch size (*B* =100), thereby capturing *K* types of useful patterns from the regulatory features of candidate SNPs prior to training. If there are 100 chromosomal blocks and each block contains 30 candidate SNPs, the total number of SNPs for pre-training is 3,000. We initialized each weight at the beginning of the pre-training as values sampled from the normal distribution, $N(0, \sqrt{6/(M+K)})$, where *M* is the number of features of a SNP and *K* is the number of hidden nodes. The total number of pre-training epochs was set to 200. All algorithms were implemented using the Theano library (7).

#### Hyperparameters

Deep neural networks rely on a large number of hyperparameters, which affect their performance substantially. While testing performance on a total of 188,160 hyperparameter sets (Supplementary Fig. 2), we observed that model performance did not considerably improve with more than 100 hyperparameter sets. Therefore, we used 500 randomly sampled hyperparameter sets for reasonable computation time. Specifically, the hyperparameters that were used included the corruption level ($\alpha$), training learning rate ($\varepsilon$), momentum ($\mu$), and $\lambda_p$ for $L_p$ regularization. The range of values used for each hyperparameter is detailed in Supplementary Fig. 2. The best hyperparameter set was selected by the *F1* score on the validation data set.

#### Supplementary Methods S2: Functional analyses of predicted variants

We performed the following analyses to assess the biological function of the predicted causal variants.

##### TF binding and allelic imbalance analysis

Accurate identification of TF-contacting sequences has been made possible through the nucleotide-resolution analysis of DNase I cleavage patterns (8). We obtained these TF footprint data previously generated in 41 cell types by the ENCODE Project (http://hgdownload.cse.ucsc.edu/goldenPath/hg19/encodeDCC/wgEncodeUwDgf/files.txt). The median size of the footprints was 9 bp. We screened all heterozygous variants from the BAM files for DNase footprint sequencing. For more accurate variant detection, we carried out several clean-up procedures, including duplicate removal by the Picard tools (http://broadinstitute.github.io/picard), and performed local realignment and base quality recalibration using the Genome Analysis Tool Kit (GATK) (9). After variant calling, GATK variant filtration was performed to retain the sites for which the map quality (MQ) was ≥ 30 and the Phred scaled probability that a polymorphism exists (QUAL) was ≥ 30. We used heterozygous variants with a minimum read depth of 8 in the footprint data and obtained the degree of allelic imbalance as the log2 ratio of the number of reads carrying each allele. We tested whether the ratio of reads supporting each allele (allele 1:allele 2) deviated from 50:50 by using the binomial test (10). To correct the type I error for multiple testing, we performed the Benjamini-Hochberg correction method at the P value threshold of 0.05 (11). We then compared the fraction of the significantly imbalanced variants between positive and negative calls.

##### Evolutionary conservation analysis

The PhastCons scores represent the posterior probability that the given site is evolutionarily conserved (12). The scores for primates, placental mammals, and vertebrates are available from ftp://hgdownload.cse.ucsc.edu/goldenPath/hg19/phastCons46way/. We computed the odds ratio (OR) of positive calls in conserved regions to their odds in non-conserved regions as follows:

$$\mathrm{OR} = \frac{\dfrac{\text{positive SNPs in conserved regions}}{\text{negative SNPs in conserved regions}}}{\dfrac{\text{positive SNPs in non-conserved regions}}{\text{negative SNPs in non-conserved regions}}}.$$

For conserved regions, we searched association blocks for DNA sequences with the primate PhastCons score > 0.5. The non-conserved regions were defined as having the PhastCons score < 0.5. Positive and negative SNPs in all association blocks were counted. In addition, we compared the mean PhastCons score at positive and negative calls.

#### Target gene function analysis

We performed gene set enrichment analysis to determine the function of genes that are targeted by putative causal SNPs. Target genes were annotated by DHS correlation and long-range chromatin interaction data for psychiatric disorders and autoimmune diseases, respectively. As for psychiatric disorders, we assigned the gene as a target gene for the given SNP when a DHS that includes a causal SNP was highly correlated with a DHS that is included in a gene body or gene promoter (5kb upstream of the transcription start site (TSS)) across a large number of cell types. The DHS correlation data was obtained from a previous publication, where a high DHS correlation was defined as Pearson correlation coefficient r ≥ 0.7 between two DHSs which were 500kb apart at maximum (13). As a result, a total of 353 target genes were identified for 1,283 ADHD SNPs. Likewise, 426, 466, 371, and 821 genes were identified for 1,277 ASD SNPs, 2,090 BPD SNPs, 1,424 MDD SNPs, and 2,462 SCZ SNPs, respectively. Genes at the major histocompatibility complex locus were excluded from the analysis. Because of high gene density in this region, it is unlikely to improve the power of enrichment test in most cases (14). Regarding autoimmune diseases, we assigned a gene to a given SNP when the gene promoter (from 1.5kb upstream of the TSS to 0.5kb downstream of TSS) has connection with an enhancer that includes the SNP based on long-range interaction data.

As a result, 1,439, 1,931, 1,793, and 1,732 genes were identified for 4,113 RA SNPs, 4,358 SLE SNPs, 5,801 CD SNPs, and 5,020 SNPs, respectively. We employed seven chromatin interactome datasets (15–18) (listed in Supplementary Table 5) that were derived from different technologies encompassing chromatin interaction analysis by paired-end tag (ChIA-PET) sequencing, capture Hi-C, and integrated methods for predicting enhancer targets (IM-PET) in lymphoblastoid, K562, Jurkat, and CD4 T cells. We only used intrachromosomal interactions. We then conducted gene set enrichment analysis for target genes of each disorder using Enrichr, a freely available online tool (19). Among many available gene-set libraries, KEGG, Reactome, Panther, GO Biological Process, GO Cellular Component, dbGaP, and Disease Perturbations from GEO were employed. Next, we generated a list of search keywords (autism, autistic, attention, bipolar, depress, schizophrenia, alcohol, addiction, Parkinson, Alzheimer, axon, neuro, nervo, synap, circadian, behavior, mental, brain, calcium, cholin, adren, glutamate, dopamine, seroton, GABA, nicotine and oxytocin) that can be used for searching functional terms related with psychiatric disorders. For autoimmune diseases, we used other keywords (immune, inflammatory, cytokine, chemokine, T cell, B cell, lympho, NK T cell, natural killer cell, leukocyte, granulocyte, interferon, transforming growth factor beta, interleukin, rheumatoid, lupus) for searching functional terms. When determining the significance level, adjustment for multiple testing was performed as previously suggested (11). We eliminated terms in which >50% of the genes were originated from a single SNP.
